# Structural dynamics of DNA strand break sensing by PARP-1 at a single-molecule level

**DOI:** 10.1101/2022.02.26.482089

**Authors:** Anna Sefer, Eleni Kallis, Tobias Eilert, Carlheinz Röcker, David Neuhaus, Sebastian Eustermann, Jens Michaelis

## Abstract

Single stranded breaks (SSBs) are the most frequent DNA lesions threatening genomic integrity. A highly kinked DNA structure in complex with human PARP-1 led to the proposal that SSB sensing in Eukaryotes relies on dynamics of both the broken DNA double helix and PARP-1’s multi-domain organization. Here, we directly probe this fundamental yet poorly understood process at the single-molecule level. Quantitative SM-FRET and structural ensemble calculations reveal how PARP-1 binding converts DNA SSBs from a largely unperturbed conformation into the highly kinked state. Binding of the second N-terminal zinc finger yields an intermediate DNA conformation that can be recognized by the first zinc finger. Thus, the data is inconsistent with a conformational selection model, instead an induced fit mechanism via a multi-domain assembly cascade drives SSBs sensing. Interestingly, a clinically used PARP-1 inhibitor niraparib shifts the equilibrium towards the unkinked state, whereas the inhibitor EB47 stabilizes the kinked state.

## Introduction

Accurate sensing of DNA lesions is essential for genome maintenance across all kingdoms of life. High resolution structures of DNA lesions in complex with sensor proteins provided key insights into the underlying mechanism of damage recognition. Notably, many of these structures exhibit a highly perturbed DNA conformation that differs not only from B-form geometry but also from their respective unbound conformation. A prominent example is the binding of a G-T mismatch by the mismatch repair protein MutS, which causes the damaged DNA to adopt a kinked state (Lamers et al. 2000). Similarly, a nicked DNA bound by the Flap endonuclease Fen1 is sharply kinked at the damage site (Tsutakawa et al. 2011). DNA kinking is also observed upon Rad4/XPC binding to damaged DNA (Chen et al. 2015). Thus, altered DNA deformability likely plays a major role for DNA damage recognition.

In order to unravel the role of such perturbations in DNA damage recognition, single molecule studies have been employed. Single molecule Förster resonance energy transfer (smFRET) data provided further insight into DNA kinking upon binding of MutS protein, showing that the DNA fluctuates rapidly between different bent states (Sass et al. 2010). DNA conformational dynamics are also important during Fen1 binding to a DNA flap, which results in a change in DNA kinking that in turn leads to flap junction opening (Sobhy et al. 2013; Craggs et al. 2014; Rashid et al. 2017). Remarkably, conformational changes of DNA during DNA damage recognition are also observed in the context of chromatin since the UV-damaged DNA-binding (UV-DDB) protein complex accesses inward lesions on the nucleosome by shifting the nucleosomal DNA (Matsumoto et al. 2019). All these examples show that perturbations in DNA conformation often play a key role in DNA damage recognition. However, as yet no mechanistic insight was obtained about the pathway by which the protein-DNA complex reaches this state leaving key determinants of DNA damage recognition elusive.

Understanding the mechanistic basis of DNA damage sensing is particularily relevant for Poly(ADP-ribose)Polymerase-1 (PARP-1). PARP-1 is a highly abundant and conserved nuclear stress response protein and constitutes the principal sensor of DNA single stranded breaks and other types of DNA lesions in higher Eukaryotes. Moreover, it is also involved in a plethora of other nuclear processes including transcription regulation as well as chromatin organisation (Gupte, Liu, and Kraus 2017; Ray Chaudhuri and Nussenzweig 2017; Hanzlikova et al. 2018; Rudolph et al. 2018; Wang, Luo, and Wang 2019). Upon binding to DNA damage and by other signals, PARP-1 becomes allosterically activated and catalyzes a post-translational modification, in which poly(ADP-ribose) (PAR) is synthesized and attached to the sidechain of proteins, thus directly modulating their function and leading to the recruitment of various downstream factors (Wei and Yu 2016). Inhibition of PARP-1 is synthetically lethal for cells deficient in DNA repair factors (e.g. BRCA1) and other stress response proteins. Such deficiencies are frequently caused by cancer mutations. Therefore, PARP-1 inhibitors are successfully used in cancer therapy and serve as a paradigm for the development of a novel generation of drugs exploiting the principle of synthetic lethality (Lord and Ashworth 2017; Ohmoto and Yachida 2017; Zandarashvili et al. 2020; Slade and Eustermann 2020).

The dynamic multi-domain architecture of PARP-1 is thought to play a central role for DNA damage recognition, for allosteric activation as well as for the design and function of PARP-1 inhibitors. PARP-1 consists of six domains (Fig. 1a), namely two N-terminal zinc fingers (F1 and F2), a third (structurally unrelated) zinc finger (F3), a BRCT-domain (breast cancer susceptibility protein C-terminal domain) containing auto-modification sites, a WGR-domain, and the C-terminal catalytic domain. The catalytic domain comprises a regulatory helical subdomain (HD) and the ADP-ribosyl transferase subdomain (ART). When free in solution, these six domains are largely non-interacting, like “beads-on-a-string” (Lilyestrom et al. 2010), but once F1 and F2 recognize a DNA lesion all domains, apart from the BRCT domain, assemble into a well-defined structure at the damage site by forming interdomain contacts (M.-F. Langelier et al. 2012; Mansoorabadi et al. 2014; Eustermann et al. 2015; Steffen, McCauley, and Pascal 2016). This multi-domain assembly cascade causes local destabilization of the HD subdomain, which releases the autoinhibitory effect of this subdomain and allows the protein to reach its catalytically active state (M. Langelier et al. 2012; Dawicki-McKenna et al. 2015; Steffen, McCauley, and Pascal 2016; M. F. Langelier et al. 2018). The importance of PARP-1’s dynamic nature is also underlined by the fact that the lethal effect of PARP-1 inhibitors is thought not to be primarily due to a lack of PAR signaling but due to the ability of the inhibitor to trap the enzyme at the DNA damage site by stabilizing the chain or inter-domain interaction (Murai et al. 2012; Zandarashvili et al. 2020).

**Fig. 1.**
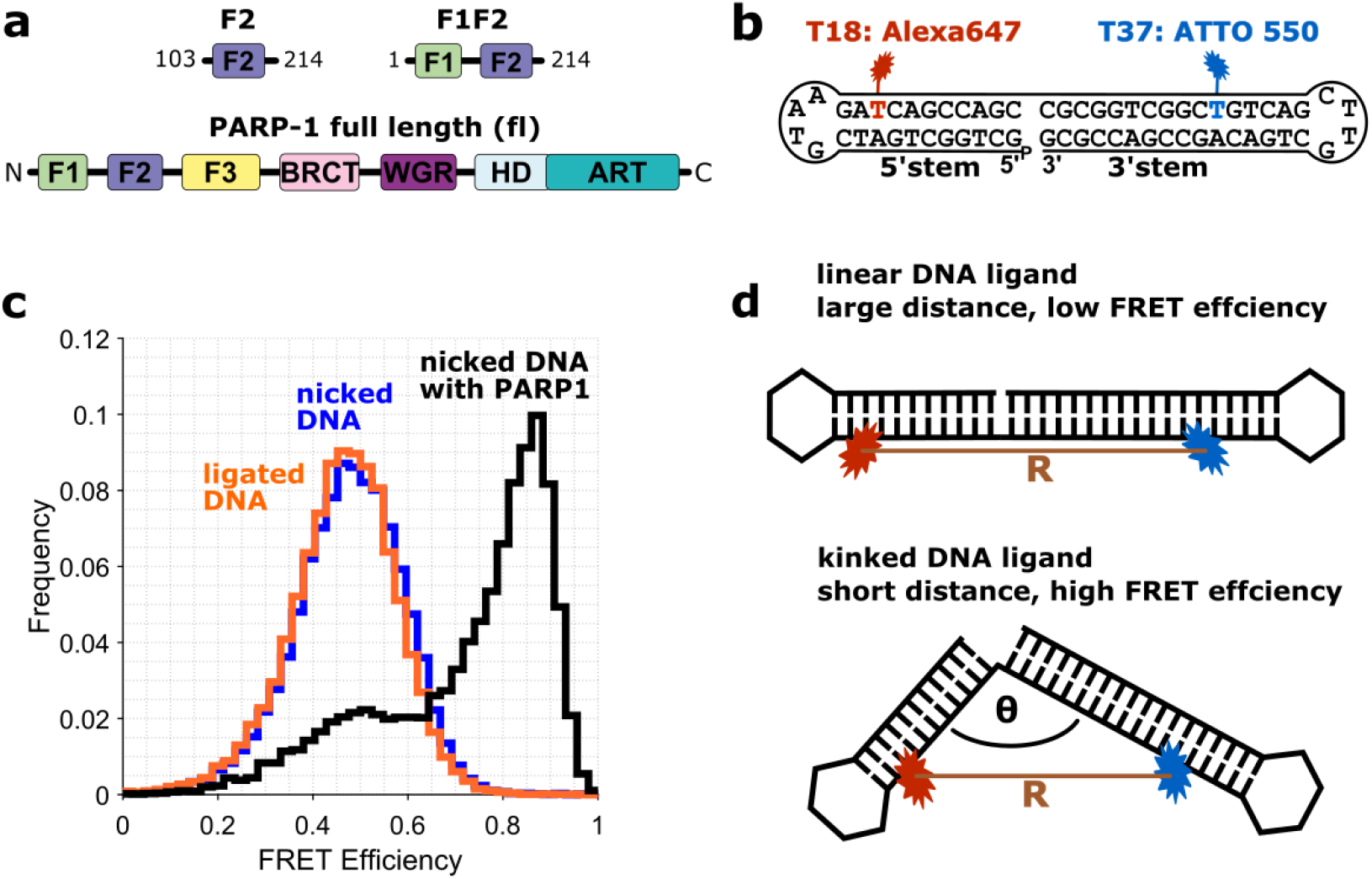
smFRET assay for probing the DNA conformation in presence of PARP-1. **a**: Domain architecture of full-length PARP-1 and PARP-1 fragments F2 and F1F2. Domains are abbreviated as: F1, F2, F3: zinc finger domains; BRCT: breast cancer susceptibility protein C-terminal domain; WGR: WGR domain; HD: α-helical subdomain; ART: ADP-ribosyl transferase subdomain. **b**: Schematic displaying the sequence and predicted secondary structure of the DNA ligand used in this study. The DNA is labeled with Alexa647 (red) at position T18 and with ATTO 550 (blue) at position T37; both dyes are attached via a C2 linker (Methods). **c**: smFRET efficiency histogram obtained from freely diffusing molecules of the nicked DNA (blue), DNA after highly efficient ligation of the nick (orange, Supplementary Fig. S1) and of the nicked DNA in presence of full-length PARP-1 (black). **d**: Schematic depicting how changes in DNA conformation can be detected by smFRET.

The structure of F1F2 bound to a 1nt gap DNA (Eustermann et al. 2015) currently remains the only high resolution structure of any PARP-1 domain in complex with an SSB, although other forms of SSBs are also known to activate PARP-1, e.g. nicks or longer gaps (Bryant et al. 2009; M. F. Langelier, Riccio, and Pascal 2014). In this structure, the gapped DNA adapts a highly kinked conformation, F2 is bound to the 3’ side of the gap, F1 binds to the 5’ side and the two zinc fingers form interdomain contacts with one another. Notably, the two flexibly linked zinc finger domains show no interdomain interaction in absence of DNA. Consequently, kinking of the DNA cannot be explained by binding to a relatively rigid, pre-formed protein interface, as proposed for other DNA damage sensor proteins (see above). Instead, it points towards a more elaborate mechanism in which altering the dynamics of the damaged DNA as well as the multi-domain organization of PARP-1 might be important for DNA damage recognition. It has been speculated that F2 is needed for the first step of SSB recognition, involving opening of the DNA around the SSB to make the F1 binding site accessible (Eustermann et al. 2011, 2015).

Many central questions regarding the mechanism of PARP-1’s damage recognition remain unanswered: does the mechanism of DNA damage recognition resemble a conformational selection or rather an induced fit, how does the behaviour of the dynamic multi-domain structure contribute to this process, and do PARP inhibitors modulate DNA damage recognition through allosteric effects? In this study, we combined smFRET experiments, structural modelling and computational approaches to gain direct insight into the binding of PARP-1 to a nicked DNA. Our data show that in the absence of protein, the DNA molecule adopts a conformational ensemble almost identical to linear undamaged DNA. We further show that when the F2 domain alone binds, it kinks the DNA, while binding of F1F2 kinks the DNA to a greater extent. By implementing a hybrid smFRET and computational approach, we determined the kinking angle of the DNA in the contexts of the different complexes. Time-resolved single molecule fluorescence also sheds light on the dynamics in these complexes and provides direct evidence that PARP-1 binding does not involve conformational selection, but instead rather resembles an induced fit mechanism. Furthermore, the functional importance of PARP-1 dynamics is supported by smFRET experiments in the presence of PARP-1 inhibitors (PARPi) such as niraparib and EB-47.

## Results

To dissect DNA SSB recognition by PARP-1 at the single-molecule level, we used a single stranded DNA molecule designed to form a dumbbell like structure, in which an SSB is created between two hairpins (in the following, the stem carrying the free 5’ terminus is called the 5’ stem and that carrying the free 3’ terminus is called the 3’ stem) (Fig. 1b, Methods). The positions of two fluorophores were optimized on either side of the nick, ATTO 550 on the 3’ stem and Alexa647 on the 5’ stem, respectively, to sensitively monitor DNA conformations by measuring the smFRET efficiency. Using time-resolved fluorescence spectroscopy (Methods), smFRET efficiencies were determined from free DNA as well as from the DNA in presence of PARP-1 (Fig. 1c). Due to the design of the DNA ligand, smFRET data can be used to assess the kinking angle between the two DNA stems (θ in Fig. 1d). Comparison of the smFRET data obtained from the DNA containing the SSB to that of a ds-DNA molecule obtained by ligation with T4 DNA ligase yielded virtually identical smFRET efficiency histograms (Fig. 1c), with peak FRET efficiencies of E=0.50±0.03 and of E=0.48±0.03, respectively (Supplementary Fig. S2a,b and Supplementary Table S1). Apparently, under the buffer conditions used here, the nicked DNA adopts a linear conformation in solution which is stabilized by stacking interactions, similarly to that of normal B-DNA.

### PARP-1 binding leads to a pronounced kink in the DNA ligand containing a SSB

Binding of PARP-1 introduces a pronounced kink in the DNA and therefore leads to a shift of the observed smFRET efficiency histogram to higher values (high FRET peak at E_HF_=0.87, Fig. 1c, Supplementary Fig. S2e). Moreover, the observed histogram is asymmetric with a pronounced shoulder at lower FRET efficiencies, indicating conformational heterogeneity of the DNA-protein complex. Interestingly, when using a fragment of PARP-1 containing only its first two zinc finger domains (residues 1-214, F1F2), we observe a smFRET histogram practically indistinguishable from that in presence of the full length protein (Fig. 2, green E_HF_=0.86 see also Supplementary Fig. S2d, E and Supplementary Table 1). Thus, binding of the first two zinc fingers is sufficient for forming the fully kinked DNA structure. In contrast, when using a shorter fragment of PARP-1 containing only zinc finger domain 2 (residues 103-214, F2) the recorded smFRET data yielded a FRET efficiency histogram shifted towards higher efficiencies as compared to DNA alone, but the efficiencies are not as high as those observed in the presence of F1F2 (Fig. 2). The histogram for the F2 complex appears comparatively broad and again not entirely symmetric. This could be explained either by a remaining fraction of unbound DNA, by structural heterogeneity of the bound DNA complex, or by conformational changes on a time-scale faster than the experimental observation time (about 1ms). Taken together, these results show that F2 alone is able to induce a kink in the DNA, but kinking is not as pronounced as that caused by binding of F1F2.

**Fig. 2.**
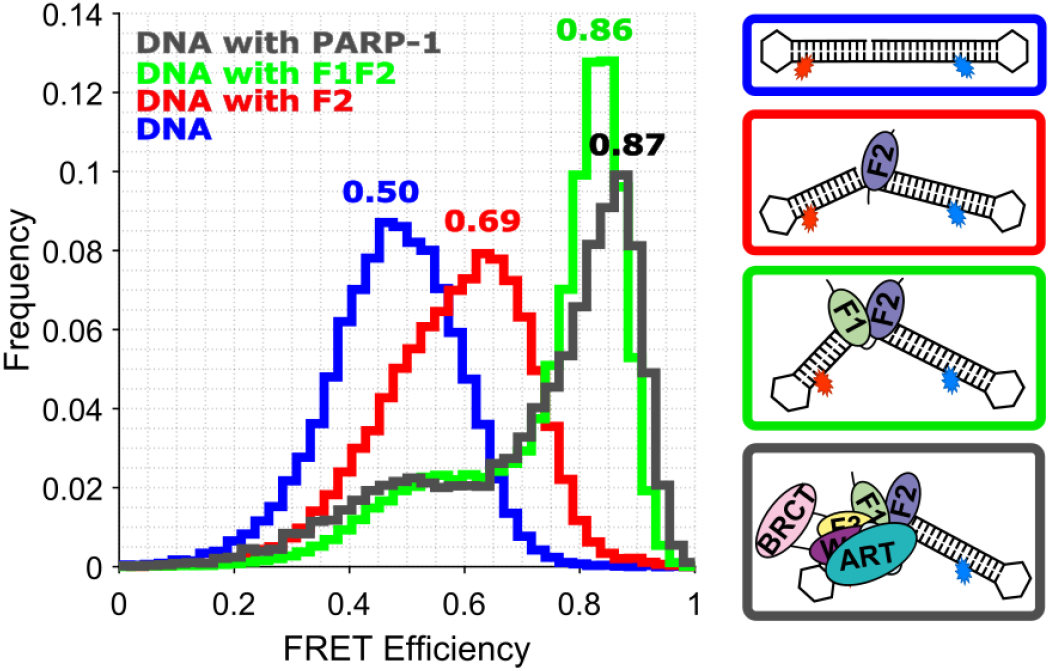
DNA conformation during nick recognition probed by smFRET. smFRET efficiency histogram of nicked DNA (blue) and nicked DNA in presence of either 10 μM F2 (red), 1 μM F1F2 (green) or 1 μM full length PARP-1 (grey here, the same histogram as in Fig. 1), together with cartoons illustrating the underlying conformations. The indicated values correspond to the peak FRET efficiencies obtained by Gaussian fitting (Methods, Supplementary Fig. S2 and Supplementary Table S1).

While smFRET data provide a very sensitive tool for investigating structural changes, changes in smFRET efficiencies can also be caused by changes in the photo-physics of the dye molecules. We therefore performed control experiments with a slightly modified DNA construct as well as a different donor dye molecule. These experiments yielded closely comparable results (Methods, Supplementary Fig. S3), confirming our interpretation that the observed changes in smFRET efficiency histograms are caused by structural re-arrangement.

### Quantification of kink angles

Next, we wanted to quantify the degree of kinking of the DNA bound to different PARP-1 fragments, namely F2 and F1F2. While smFRET data contains distance information, such an interpretation is not straightforward, since the 1D distance information about the inter-dye distance cannot be directly converted into a 3D model structure. However, if a model structure is available for a particular conformation, computational approaches can be used to compute the expected smFRET efficiency (Muschielok et al. 2008; Kalinin et al. 2012; Beckers et al. 2015; Peulen, Opanasyuk, and Seidel 2017; Reinartz et al. 2018; Nagy, Eilert, and Michaelis 2018; Eilert et al. 2018). This computed efficiency can then be compared to the experimentally measured FRET efficiency in order to establish whether or not the structure is in agreement with the experimental data, taking into account error estimates for both the smFRET efficiency measurement and the smFRET efficiency computation. Here, we integrated such a “backwards” approach with structural ensemble calculations in order to identify, at a single-molecule level, structural heterogeneity DNA conformations which agree with our experimental smFRET data and to analyze the distribution of kink angles in these ensembles in order to determine the extent of DNA kinking in each complex. The workflow is illustrated in Fig. 3a and explained in the following. Simulations can predict the measured FRET efficiency for a given model structure

**Fig. 3.**
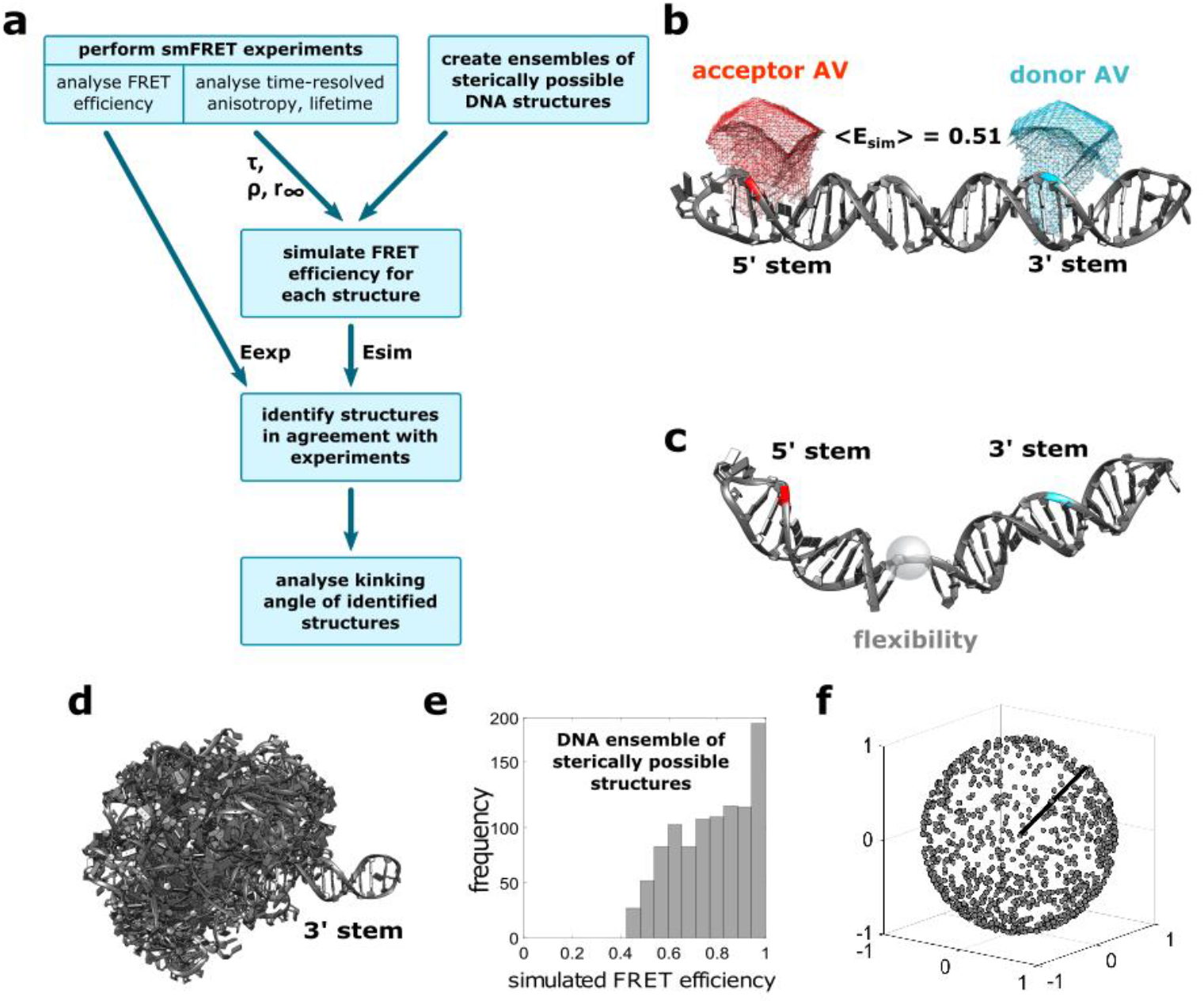
Structural analysis of DNA conformation using a computational approach. **a**: Workflow for determining the kinking angles of DNA bound to different PARP-1 fragments. For an ensemble of possible DNA conformations, the expected FRET efficiencies are simulated, taking experimental data into account, i.e. Förser radius, fluorescence lifetime and time-resolved anisotropy. From the ensemble of conformations, structures are selected for which the simulated FRET efficiency is in agreement with the measured efficiency within the given uncertainties. Eexp: experimental FRET efficiency; Esim: simulated FRET efficiency; τ: lifetime; ρ: rotational correlation time; r^∞^: residual anisotropy. **b**: Model structure for the straight DNA ligand modelled in B-form. The bases to which donor and acceptor are attached are highlighted in cyan and red, respectively. The corresponding computed accessible volumes (AV) are shown by meshes in the respective colors. **c**: Example of a structure taken from the computed ensemble of sterically possible DNA structures; the bases to which the dyes are attached are highlighted in cyan (donor) and red (acceptor), respectively. The transparent sphere indicates the flexibility of the DNA backbone in the linker region connecting the two stems. **d**: Ensemble of computed model structures for the DNA ligand alone. Structures are superposed using the 3’ stem of the DNA, and the 5’ stem is allowed to adopt any sterically possible orientation relative to the 3’ stem. For visual clarity, only 100 randomly chosen structures are shown, extracted from the full set of 1000 calculated structures. **e:** Histogram of simulated FRET efficiencies for all 1000 structures of the DNA ensemble represented in D. **f:** Representation of all simulated DNA structures in a spherical coordinate system. All structures are aligned with respect to the 3’ stem. The axis of the 3’ stem is represented by a black line. The grey dots indicate the position of the tip of the 5’ stem for each DNA structure.

In order to simulate the FRET efficiency of a given structure, the first step is to calculate the accessible volume (AV) of both dyes, given the attachment points of the dyes and the geometry of the flexible linkers that attach them to the DNA (Methods) (Muschielok et al. 2008). Examples of the resulting AVs are shown for the structure of straight DNA (Fig. 3b), and represent the space that the dye can access. Once the AVs are calculated, we use Markov Chain Monte Carlo simulations together with Bayesian Parameter Estimation (Eilert et al. 2018) to determine the expected FRET efficiency for a given structure (Methods). We first tested this approach using a model structure of the DNA in a linear, i.e. B-form, DNA conformation (Fig. 3b). The simulation yields a predicted FRET efficiency for this structure of Esim=0.51 ± 0.04 which agrees well with the measured value of Eexp=0.50 ± 0.03. The given error for the simulated FRET efficiency arises predominantly from the uncertainty in the measured isotropic Förster radius of 3% (R_iso_ = 70 Å ± 2 Å, Methods), which translates to an uncertainty of ΔE=0.04 for the simulated FRET efficiency.

### Simulations can predict the measured FRET efficiency for a given model structure

In order to simulate the FRET efficiency of a given structure, the first step is to calculate the accessible volume (AV) of both dyes, given the attachment points of the dyes and the geometry of the flexible linkers that attach them to the DNA (Methods) (Muschielok et al. 2008). Examples of the resulting AVs are shown for the structure of straight DNA (Fig. 3b), and represent the space that the dye can access. Once the AVs are calculated, we use Markov Chain Monte Carlo simulations together with Bayesian Parameter Estimation (Eilert et al. 2018) to determine the expected FRET efficiency for a given structure (Methods). We first tested this approach using a model structure of the DNA in a linear, i.e. B-form, DNA conformation (Fig. 3b). The simulation yields a predicted FRET efficiency for this structure of Esim=0.51 ± 0.04 which agrees well with the measured value of Eexp=0.50 ± 0.03. The given error for the simulated FRET efficiency arises predominantly from the uncertainty in the measured isotropic Förster radius of 3% (Riso = 70 Å ± 2 Å, Methods), which translates to an uncertainty of ΔE=0.04 for the simulated FRET efficiency.

The next step towards determining the kinking angle of the nicked DNA molecule was to generate a large ensemble of sterically possible DNA conformations by keeping both stems in fixed conformations while varying their relative orientation (with the respect to the flexible region at the nick, Fig. 3c) in any way that did not lead to steric clashes between the stems (Methods). This leads to a very heterogeneous distribution of sterically allowed DNA conformations (Fig. 3d). For each of these conformation we then simulated the expected FRET efficiency. The resulting simulated FRET efficiencies show a broad distribution covering the range from Esim=0.42 to Esim=1 (Fig. 3e). Fig. 3f illustrates these very DNA structures in a spherical coordinate system where the angle between the two stems (3’ stem represented by the black line and 5’ stem represented by the grey dot) was computed for every structure (Methods).

Next, in order to compare the generated structural ensembles to the recorded smFRET, we selected those structures from the ensemble which agreed with the respective measured FRET efficiencies. To this end, for each structure we tested whether the simulated FRET efficiency was within ΔE=0.05 of the measured smFRET efficiency, where the uncertainty was obtained using error propagation, considering an error contribution of 0.03 from the experimental FRET efficiency and 0.04 from the simulated FRET efficiency (Methods). From the ensemble of 1000 structures, 87 structures were in agreement with the smFRET data for DNA alone (i.e. Esim=0.50±0.05, blue structures in Fig. 4a), 154 with the data for DNA+F2 (Esim=0.69±0.05, red structures in Fig. 4a) and 199 structures with the data for DNA+F1F2 (Esim=0.86±0.05, green structures in Fig. 4a). The respective ensembles of selected structures in Fig. 4a show that the three different FRET efficiency ranges lead to distinct distributions of kink angles θ (Fig.4b and c). For a better overview of the selected structures we chose a representation which displays the respective structures with reduced complexity, highlighting only the relative directions of the two DNA stems (Methods, Fig. 4b). The side view shows that the allowed kinking angle regimes are well separated for the three ensembles (upper plot in Fig. 4c). From the view along the 3’ axis (lower plot in Fig. 4b), it is apparent that the selected structures lie on circles around the 3’ axis, indicating that our approach gives no information on the angle ϕ through which the 5’ stem is rotated. This is not surprising, as the structure selection is based on a single distance between the two dyes. From the histograms we determined the mean kinking angles of <θ>=176°±24° for DNA only, <θ>=130°±20° for DNA+F2 and <θ>=90°±14° for DNA+F1F2 (Methods). This kinking angle determination relies on the structural ensemble of the nicked DNA molecules. However, as a control we also computed an alternative structural ensemble by modeling F2 bound to the 3’ stem (Methods) and again simulated the expected FRET efficiency for each structure in the ensemble (Supplementary Fig. S4). Again we compared the simulated FRET efficiencies to the experimental data given the uncertainty of ΔE=0.05 to define structural sub-ensembles for the complex in presence of F2 or F1F2 which are in agreement with the experimental data (Supplementary Fig. S4). From the respective structural ensembles, we again computed the mean kinking angles <θ>=127°±19° for DNA+F2 and <θ>=87°±15° for DNA+F1F2, which is in excellent agreement with the analysis based on the structural ensemble of the DNA only.

**Fig. 4.**
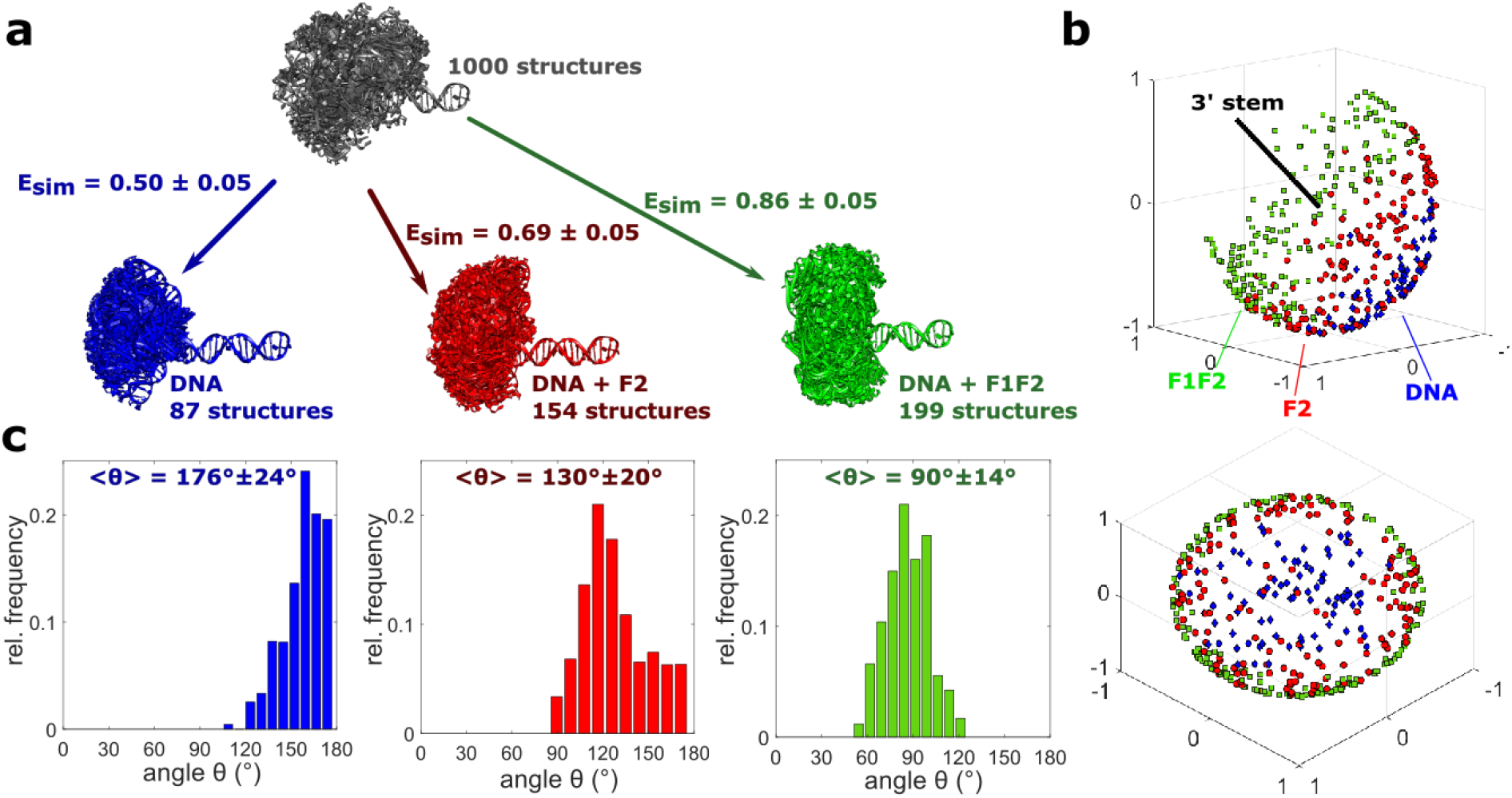
From structural ensembles to kinking angle histograms. **a:** An ensemble of 1000 possible structures for the DNA was calculated (only 100 structures are shown for clarity, grey). From this ensemble, structures in agreement with the experimental FRET efficiencies were selected, i.e. 87 for free DNA (blue), 154 for DNA+F2 (red) and 199 for DNA+F1F2 (green). **b**: Combined representation of the selected structures shown in a. All structures are aligned with respect to the 3’stem. The axis of the 3’ stem is represented by a black line. The colored objects indicate the position of the tip of the 5’ stem for each selected structure (blue diamonds for free DNA; red circles for DNA+F2; green squares for DNA+F1F2). Two different views of the 3D plot are shown, a side view (top) and a view along the 3’ axis (bottom). **c**: Kinking angle distributions for the selected structures shown in a. The mean and standard deviation of the angles are indicated for each histogram.

### Dynamic analysis reveals fast dynamics for nick DNA in presence of PARP-1

As described above, the observed smFRET histograms for DNA in presence of F2, F1F2 and full length PARP-1 showed shoulders at lower FRET efficiencies (Fig. 2). A possible reason for this could be averaging over different structural states that interconvert on a time scale on the order of 1ms. We therefore used several analytical approaches to test the data for evidence of dynamics in these structures (Supplementary methods). First, we investigated the dependence of smFRET efficiency on burstwise donor lifetimes in presence of the acceptor (Supplementary Fig. S5a). When comparing these lifetimes with the ratiometrically determined FRET efficiencies, one can analyze in two-dimensional histograms whether the observed histograms fall on the so called static FRET line or whether deviations can be seen that are caused by dynamic interconversion of linear and kinked conformations (Methods, Kalinin et al. 2010). While for the free DNA the observed 2D histogram falls on the static FRET line (Supplementary Fig. S5a), there are clear deviations observed for the DNA in presence of F2, F1F2 or PARP-1, indicating the presence of a dynamic interconversion. Similarly, burst variance analysis (BVA, Torella et al. 2011) also showed indications of dynamic interconversion for DNA+F2, DNA+F1F2 and DNA+PARP-1, but not for the DNA in absence of proteins (Supplementary Fig. S5b). More direct information about the respective dynamics with time scales between 0.2 and 5ms comes from time-window analysis, where histograms of FRET events were computed using different time-binnings (Methods, Supplementary Fig. S5c). Again, for the case of DNA alone, no dynamics were observed. Interestingly, for the DNA in presence of protein only very small changes were observed, indicating that dynamics occur on an even faster time-scale.

Besides actual dynamics, dye-photophysics could also influence the observed smFRET efficiency histogram. However, by using a Pulsed Interleaved Excitation scheme (Müller et al. 2005) we were able to investigate these effects directly. To this end, we performed FRET-FCS experiments for the single molecule burst data sub-ensemble consisting of the acceptor only species (Supplementary Fig. S6, Supplementary methods). This analysis revealed that the acceptor, Alexa647, attached to the nick DNA in the absence or in the presence of either F2 or F1F2 has a short relaxation time of 6-7 μs (Supplementary Fig. S6b, τ_phot_). This is in good agreement with previous reports (Vandenberk et al. 2018; Baibakov and Wenger 2018), and is presumably caused by cis-trans photo-isomerization, characteristic of cyanine dyes (Widengren, Mets, and Rigler 1995; Widengren and Schwille 2000). For the full length PARP-1, the acceptor relaxation time is slightly longer (∼10μs) in comparison to results for the zinc finger fragments (Supplementary Fig. S6b, τ_phot_), which is most probably due to direct interactions with the additional domains in a so called PIFE effect (Stennett et al. 2015).

To provide more evidence concerning the nature of the species involved in the dynamic interconversion between the linear and kinked DNA conformations, we employed FRET-FCS experiments (Schwille, Meyer-Almes, and Rigler 1997; Torres and Levitus 2007 Supplementary methods). These show that the translational diffusion times τ_D_ (Supplementary Fig. S7) in our confocal microscope for the low FRET population of DNA with PARP-1 (1990 μs) and for that of DNA with PARP-1 bound to niraparib (1976 μs), are clearly distinct from that of DNA alone (1303 μs). These diffusion times provide strong support for our interpretation that the protein remains bound to the DNA in these low-population states where the DNA is linear.

Next, we performed a quantitative analysis of the observed fluorescence dynamics caused by structural changes using filtered FCS (fFCS, Methods, Felekyan et al. 2012; Barth et al. 2018). From the observed smFRET efficiency histograms of both DNA+F2 and DNA+F1F2, we selected smFRET efficiency regions towards the edge of the histogram for interconversion analysis (Fig.5, blue and violet areas in the FRET efficiency distributions). By globally fitting the fFCS auto- and cross-correlation functions, the relaxation time for interconversion between the two respective species can be determined (Methods, Fig.5 and Supp. Table S2). We find relaxation times of *τ*_*F*2_ = 404 ± 62μ*s* for DNA+F2, *τ*_*F*1*F*2_ = 265 ± 26μ*s* for DNA+F1F2 and *τ*_*PARP*−1_ = 273 ± 19μ*s* for DNA+PARP-1 (Supp. Table S2), which account for the fast dynamics in the respective systems. The relaxation time is given in each case by the inverse sum of the transition rates between the two states (*τ* = (*k*_12_ + *k*_21_)^−1^), and therefore when comparing the relaxation times for DNA+F2 and DNA+F1F2 we find (*k*_12_ + *k*_21_)_*F*2_ < (*k*_12_ + *k*_21_)_*F*1*F*2_. In summary, we observe dynamics for the DNA in presence of F2, of F1F2 and of full length PARP-1, and we are able to quantify the time scale of these dynamics using fFCS.

**Fig. 5.**
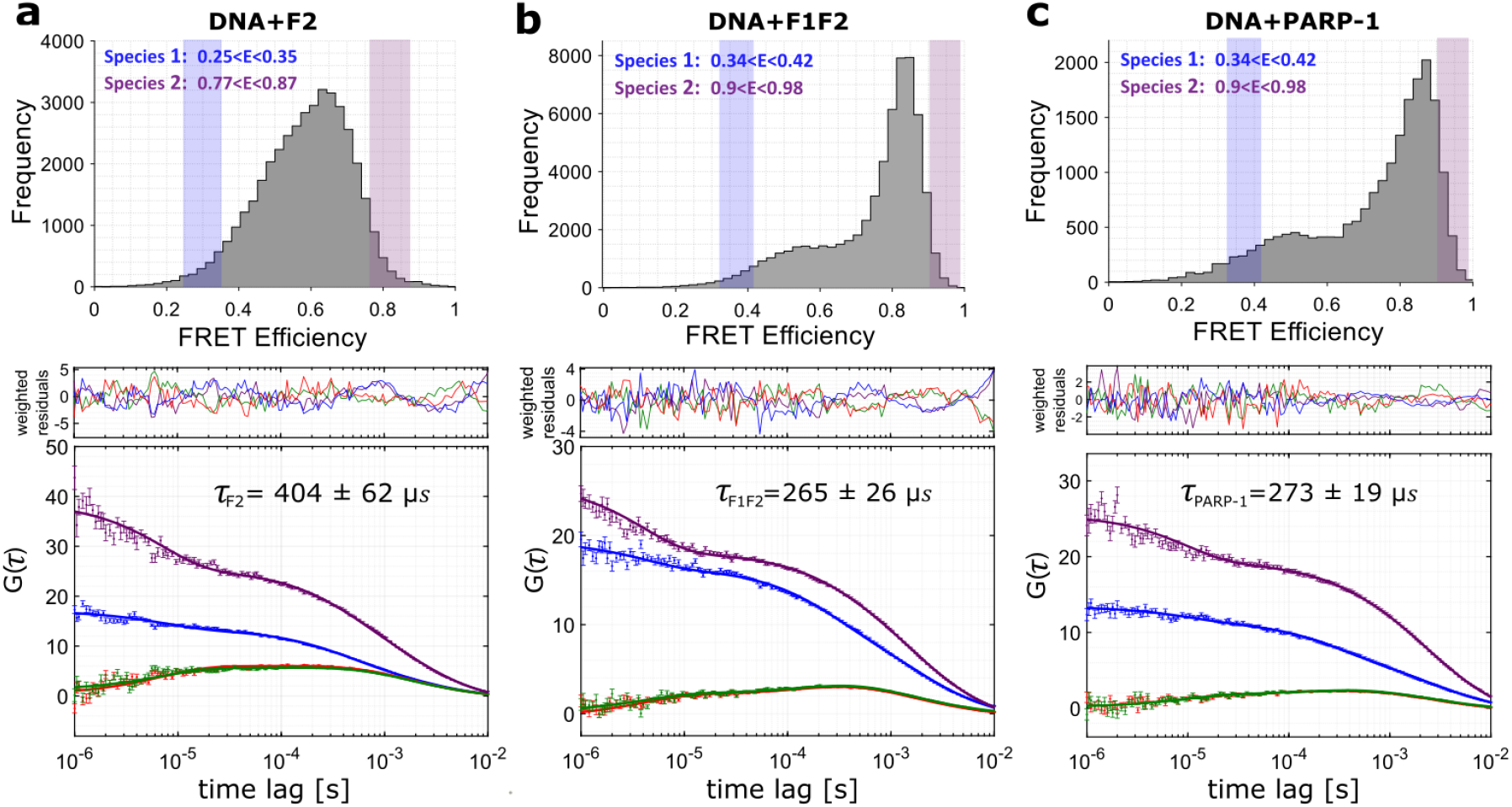
Results of fFCS analysis for DNA in presence of F2, F1F2 and full length PARP-1. Dynamic analysis using fFCS for DNA in presence of F2 (**a**), in presence of F1F2 (**a**) and in presence of full-length PARP-1 (**c**). FRET efficiency histograms with the thresholds that define species 1 (shaded blue region) and species 2 (shaded violet region) are shown in the upper panels and the corresponding fFCS fits for the auto- (violet, blue) and cross-correlation functions (green, red) in the lower panels. The middle panels show the weighted residuals for every fit of the respective correlation functions; in the top right corner of the fFCS plot the resulting relaxation times are displayed.

### PARP-1 inhibitors can influence the dynamics of PARP-1

Next, we investigated whether the observed distribution of binding states can be modulated with PARP-1 inhibitors (PARPi). Some PARPi were recently shown to drive the PARP-1 allostery and activate or inhibit its function (Zandarashvili et al. 2020; Ogden et al. 2021). Here, we characterize effects caused by EB-47 and niraparib, which have been interpreted as “pro-retention” and “pro-release” inhibitors, respectively (Zandarashvili et al. 2020; Slade and Eustermann 2020). We observe that EB-47 traps the PARP-1-DNA complex in the kinked state, as the ELF (low FRET effciency) population decreases substantially in comparison to the situation without inhibitor (Fig. 6a, magenta). Interestingly, niraparib has a different effect on the observed smFRET histogram, since in this case the ELF population is increased rather than decreased (Fig. 6b). Note, a slight shift of the position of the peaks in the smFRET histograms in the presence of niraparib relative to the other observed smFRET distributions is not due to differences in the conformation, but instead is caused by the high fluorescence anisotropy (Supplementary Fig. S8).

**Fig. 6.**
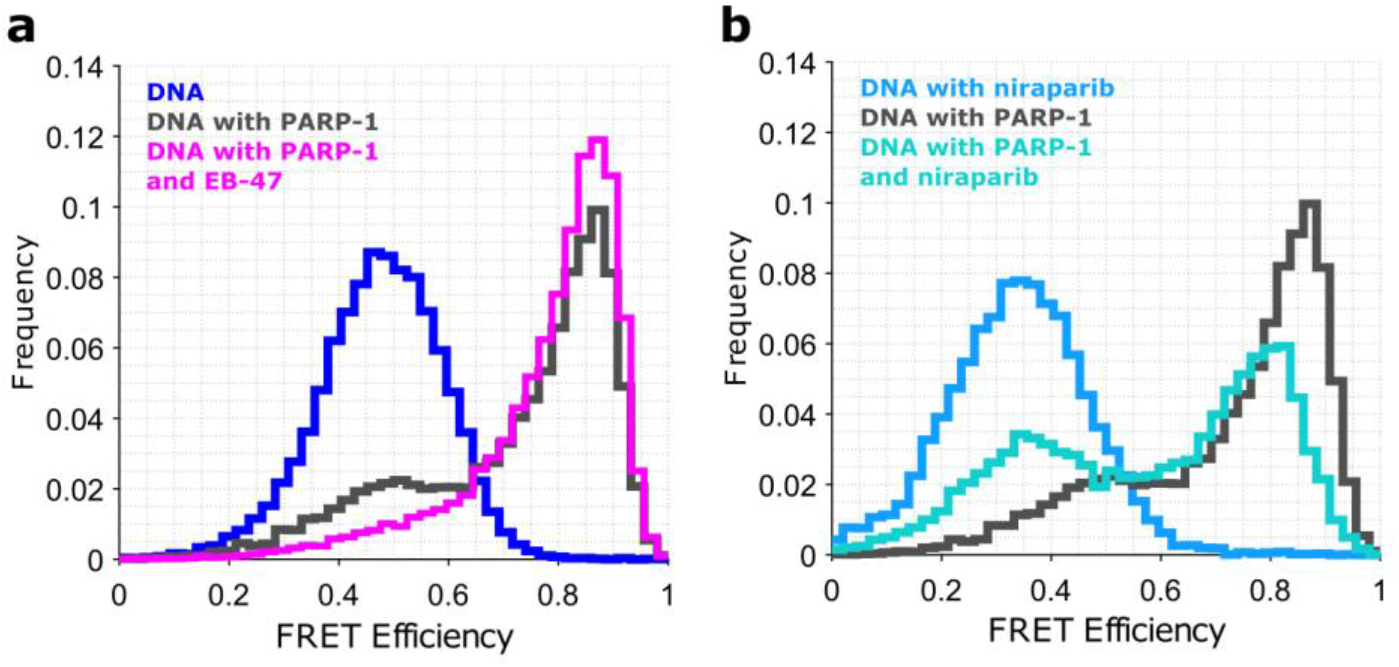
DNA conformation probed by smFRET in presence of PARP-1 and PARP inhibitors. a: smFRET efficiency histogram of nicked DNA (blue) and nicked DNA in presence of either PARP-1 (grey, same histogram as in Fig.1) or PARP-1 and EB-47 (magenta). b: smFRET efficiency histogram of nicked DNA with niraparib (light blue) and nicked DNA in presence of either PARP-1 (grey here) or PARP-1 and niraparib (turquoise);

## Discussion

In this study we have obtained quantitative, single molecule insights into the mechanism of DNA single strand breakage recognition by human PARP-1. By developing a hybrid smFRET and computational approach in conjunction with time resolved fluorescence spectroscopy analysis, we analyzed the structural heterogeneity and dynamics of nicked DNA alone, as well as when bound to full-length PARP-1 or its zinc finger domains F2 or F1F2, recapitulating the first crucial events on SBB sensing and PARP-1 activation. The free nicked DNA adopts a conformation that resembles that of intact DNA (E=0.50 for nick DNA and E=0.48 for ligated DNA Supplementary Fig. S2). It is well-known that intact DNA does not necessarily follow a strict, linear canonical B-form conformation (Wozniak et al. 2008; Wu et al. 2015). Thus, the investigated DNA template may be slightly bent, however we observe no specific effect due to the nick, and in particular no evidence for kinking at this position. This is also consistent with previous ensemble averaged NMR measurements of nicked DNA (Kozerski et al. 2001). Using the described novel hybrid single-molecule and computational approach, we determine a mean angle of <θ>=176°±24° for the free DNA. Note, that the definition of the DNA axis, i.e. the number of base pairs considered for axis calculation (Methods), has a small impact on the resulting angle between the axes, and the limited number of base pairs in each stem of the DNA leads to angles smaller than 180°, even for straight DNA (i.e. to θ=176° for our model of straight DNA). In our analysis where individual DNA structures are identified which agree with the smFRET data of DNA+F1F2, we found a mean angle of <θ>=90°±14° (when selecting from the DNA-only ensemble, Fig. 4c) and <θ>=87°±15° (when selecting from the DNA+F2 ensemble, Supplementary Fig. S4c). Taking into account the error, this result is in accord both with early electron microscopy measurements, (Le Cam et al. 1994) and with ensemble averaged NMR measurements on a gapped DNA bound by F1F2 (Eustermann et al. 2015). In order to compare our data for the nicked DNA to the published structure for the gaped DNA, we simulated F1F2 bound to the nicked DNA assuming the same interactions are formed. Applying the NMR constraints previously measured for the gapped DNA to the nicked DNA led to a very well-defined conformation, which is described by a structural ensemble of 40 structures (Supplementary Fig. S9a). From this ensemble, structures with predicted FRET efficiencies matching those of the experimental FRET data were selected. These selected structures have a mean angle between the two DNA stems of <θ>=102°±1° (Supplementary Fig. S9b), in good agreement with the data for the other two computed structural ensembles. This result suggests a quite similar conformation of the complex with either a nicked or gapped DNA. One should note that NMR measurements for F1F2 bound to a gapped DNA revealed an interaction between F1 and the unpaired base at the site of the gap itself, which is of course not present in the nicked DNA. This can lead to different kinking angles for the two complexes. Interestingly, the simulated FRET efficiencies for the ensemble of F1F2 bound to the nick yielded a mean and standard deviation of Esim=0.79±0.04 (Supplementary Fig. S9c) and 18% of the 40 structures agreed with the smFRET results (i.e. Esim=0.86±0.05), however, these are only at the upper edge of the simulated FRET efficiency histogram. Thus our data suggests that the nicked structure is kinked to an angle slightly smaller than that observed in the NMR structure of F1F2 binding to a gapped DNA. As a control, similar angles for DNA, DNA+F2 and DNA+F1F2 were obtained for the alternative DNA construct containing different dyes (Supplementary Fig. S10), showing the robustness of the method.

In recent years, large efforts have been made for using smFRET measurements as quantitative tools for obtaining structural information (Nagy, Eilert, and Michaelis 2018; Hellenkamp et al. 2018; Lerner et al. 2021). In particular, smFRET measurements using networks of measurements have provided structural information (Muschielok and Michaelis 2011; Kalinin et al. 2012; Hellenkamp et al. 2017). Here, we have shown that quantitative information from single molecule measurements of an individual distance combined with structural modeling is sufficient for obtaining quantitative structural models thus yielding direct mechanistic insight.

The smFRET data for F2 binding to a nicked DNA template provides not only structural information but also mechanistic insight into the mechanism of SSB recognition. In our analysis of DNA structures in presence of F2, we found a mean angle of <θ>=130°±20° (when selecting from the DNA ensemble, Fig. 4c) and <θ>=127°±19° (when selecting from the DNA+F2 ensemble, Supplementary Fig. S4c), which is a clearly distinct result from those for either free DNA or DNA+F1F2. Binding of F2 alone leads to a kink in the DNA that becomes significantly sharper in the context of binding by F1F2. Importantly, the sharp kinking angles of the nicked DNA template seen in the F1F2 complex are neither observed for the DNA alone, nor in presence of F2, in agreement with an induced fit mechanism for DNA damage recognition by PARP-1. Moreover, even though we do observe dynamics, results from the fFCS analysis rule out a conformational selection mechanism. The conversion time for DNA+F2 determined from the fFCS analysis (τ_F2_=404±62μs) is longer than that for DNA+F1F2, (τ_F1F2_=265±26μs), and therefore the sum of the interconversion rates is smaller. In a conformational selection model, the forward rate for formation of the partial kink by binding of 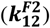 would have to be higher than or equal to that for formation of the full kink by binding of 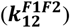, since thermal bending to an angle required for F2 binding would occur more often than thermal bending required for F1F2, where a sharper kink-angle is needed. Moreover, when comparing the smFRET histograms of DNA+F2 and DNA+F1F2 (Fig.2, Supplementary Fig. S2), the DNA+F1F2 histogram has two local maxima at about 0.6 (E_LF_) and 0.86 (E_HF_) FRET efficiency, with the higher state being the preferred state, while the F2 histogram shows only one broader peak. Given the significant difference between the two states in the DNA+F1F2 histogram, the backward rate to the linear conformation, 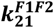, of DNA+F1F2 would have to be smaller than 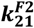 if the peak observed for F2 were due to an averaging of different states within the conformational search, in complete contrast to the fFCS data actually measured. Therefore, we can conclude that the observed state in the presence of F2 is a structurally independent state, thus ruling out the conformational selection model. Instead, protein binding leads to progressive kink formation as expected for the induced fit mechanism.

It has been suggested that binding of F2 constitutes the first crucial event not only for SSB recognition, but also for triggering the domain assembly cascade leading to PARP-1 activation (Eustermann et al. 2015). Our single molecule data provides now direct mechanistic insights into this process by showing that F2 induces DNA kinking and makes the 3’ stem binding site for F1 accessible. F2 has a higher affinity for SSBs than does F1, and isolated F1 has been shown to bind SSBs differently than it does in context of F1F2 (Eustermann et al. 2011). Presumably, both zinc fingers F1 and F2 compete for the accessible binding site on the 3’ stem of an SSB, but the higher affinity of F2 results in an F2-induced opening of the DNA that twists the two DNA stems away from each other; binding of F1 to the exposed 5’ stem as well as the interdomain interactions between F1 and F2 drives the DNA into its highly kinked conformation. This induced fit mechanism involving a multi-domain assembly and concerted DNA perturbations which differs from other DNA damage sensors for which DNA perturbations are coupled to an active process using energy derived from ATP-hydrolysis (e.g. as in the case of MutS, Lamers et al. 2000) or to conformational selection by a relatively rigid recognition surface (e.g. as in the case for the UV-damaged DNA-binding protein DDB2, Matsumoto et al. 2019). It partially resembles the induced fit described for Fen1, in which loops and helical elements of the cap become ordered upon DNA kinking by the otherwise relatively rigid recognition module (Rashid et al. 2017). The cooperative behavior of the two N-terminal zinc fingers of PARP-1 helps to explain the specificity of SSB sensing, while it is also consistent with their mutual requirement for efficient recruitment to laser induced DNA lesions in cells (Eustermann et al. 2015; Mortusewicz et al. 2007). Interestingly, the revealed pathway for an induced fit DNA damage recognition mechanism also helps to understand PARP-1 activation. F1 binding to the 3’ stem has been shown to trigger the cascade of domain-domain interactions that leads to activation of PARP-1’s C-terminal catalytic domain (M. Langelier et al. 2012; Eustermann et al. 2015), explaining previous data that F2 is needed for PARP-1 activation by an SSB, whereas it is dispensable for recognition of a DSB, where the F1 binding site is not obscured by a second stem (Ikejima et al. 1990).

Moreover, even though the amplitude of the observed high FRET peak is quite high, analysis of the observed dynamics in the smFRET data shows that even the highly kinked DNA state is not completely stable, and transient excursions to an (almost) linear DNA conformation are revealed by the fFCS analysis. In this context, we note that a linear DNA conformation of DSBs has also been observed for the related PARP-2 protein in the presence of nucleosomes (Bilokapic et al. 2020; Rudolph et al. 2021). PARP-2 contacts DSBs via its WGR domain. Given that the WGR domain of PARP-1 has been implicated in DNA intersegment transfer during damage search (Liu et al. 2017; Rudolph et al. 2018) it is conceivable that a similar configuration can be also adapted by PARP-1 in context of SSBs or intact DNA.

PARP-1 is an important clinical target in the treatment of cancer. Here, PARP inhibitors are used to trap PARP-1 at DNA breaks. Recently, using hydrogen exchange, three classes of inhibitors have been defined based on their allosteric action; while class I inhibitors such as EB-47 stabilize the bound conformation, class III inhibitors weaken the interaction with the DNA damage (Zandarashvili et al. 2020; Ogden et al. 2021). Here, our smFRET data of PARP-1 provides novel mechanistic insight. For the case of EB-47, the observed effect matches our expectations. The kinked DNA conformation is stabilized, and the low FRET state for the linear conformation virtually disappears (Fig. 6a). Interestingly, the class III PARP-1 inhibitor niraparib has the opposite effect, the kinked state is destabilized as predicted by the hydrogen exchange data (Zandarashvili et al. 2020) and the linear conformation becomes more prominent (Fig. 6b). While the observed smFRET values could indicate that niraparib causes the dissociation of PARP-1, this is not what is observed. In the highly dilute situation of the single molecule experiment, re-binding of PARP-1 would be fairly slow and one would expect the equilibrium to shift even more to the side of the low FRET peak in a PARP-1 concentration dependent manner. However, more direct evidence, that the observed low FRET population in presence of niraparib is due to a linear conformation of PARP-1 bound to the nicked DNA comes from a FRET-FCS analysis of the low FRET population (Supplementary Fig. S7). Here, the observed diffusion time of 1976μs matches that of the DNA+PARP-1 in absence of inhibitor (1990μs) and is clearly distinct from that of free DNA (1303μs), showing that the inhibitor does not lead to an increased unbinding but rather to an increase in the population of a linear DNA conformation bound to PARP-1. This also is in agreement with the observation that niraparib doesn’t change the unbinding rate of PARP-1 when bound in the context of nucelosomal substrates (Rudolph et al. 2018).

## Summary, conclusion, outlook

We used time-resolved smFRET measurements and a computational approach to investigate the role of the zinc fingers F1 and especially F2 in the recognition of DNA nicks by PARP-1. Our results show that F2 alone is able to introduce a kink at the nick, which is not present in the isolated DNA. We also show that further kinking at the nick occurs in the presence of F1F2. These findings support a model in which F2 opens the DNA at an SSB and makes the binding site for F1 accessible, but only binding of both zinc fingers together leads to the highly kinked conformation providing a possible pathway for DNA damage recognition as well as the first crucial steps for catalytic activation of PARP-1 via successive domain assembly. The quantitative analysis of the observed smFRET dynamics using fFCS allows us to rule out a conformational selection model and provides strong evidence for the induced fit mechanism. Interestingly, smFRET analysis shows the presence of an additional, minor linear DNA conformation in the bound state of PARP-1 and a dynamic interconversion between major and minor states. The equilibrium between these two structural states is altered by PARP-1 inhibitors. Future studies could address whether the linear state represents an intermediate in the unbinding pathway of PARP-1 and how other proteins of the repair machinery alter the observed dynamic equilibrium.

## Materials and Methods

### Preparation of DNA ligands

The double labeled single-stranded DNA (sequence 5’-GCTGGCTGATCGTAAGATCAGCCAGCCGCGGTCGGCTGTCAGCTTGCTGACAGCCGACCGCG-3’, with Alexa647 attached to T18 via a C2 linker and ATTO550 attached to T37 via a C2 linker, referred as DNA_Atto_, Fig. 1a) was received in two parts which were ligated to produce the final dumbbell like structure. The acceptor strand (sequence 5’-GCTGGCTGATCGTAAGATCAGCCAGCCGCGGTCG-3’ with Alexa647 attached to T18 via a C2 linker, was synthesized by IBA GmbH (Göttingen). The donor strand with sequence 5’-GCTGTCAGCTTGCTGACAGCCGACCGCG-3’ with an amino C2 modification at T26 was synthesized by Biomers (Ulm). For labeling, the lyophilized oligos were dissolved in TE buffer (10 mM Tris pH8.0, 1mM EDTA) and then mixed 1:1 with a 100mM sodium tetraborate buffer pH 8.5 to a final DNA concentration of 100 μM in 50μl. The labeling reaction was started by adding 50nmol Atto550-NHS-Ester (Sigma-Aldrich, dissolved in DMSO). After incubation for 1 hour at 37°C, another 50nmol of dye were added and incubated overnight at 37°C. The labeled DNA was then separated from free dye and unlabeled DNA by denaturing PAGE and gel extraction. The donor-labeled DNA was phosphorylated at the 5’ end using T4 Polynucleotide Kinase (New England BioLabs) for 1 hour at 37°C, followed by a heat inactivation.

For ligating the donor and acceptor strands, both were mixed (∼5μM acceptor strand and ∼2 μM phosphorylated donor strand in a final volume of 19μl) in T4 RNA ligation buffer (New England BioLabs) supplemented with 1mM ATP (New England BioLabs). The strands were annealed by heating to 90°C for 30 seconds followed by an incubation for 4 minutes at 42°C, decreasing the temperature by 1°C per minute down to 36°C and incubating again for 4 minutes. After annealing, the sample was ligated overnight at 16°C using T4 DNA ligase (New England BioLabs).

Another DNA construct with identical sequence but with Alexa647 attached to T18 via a C6 linker and 6-Tamra attached to T37 via a C6 linker (referred as DNA_Tamra_) was synthesized by IBA GmbH (Göttingen). The desalted DNA was purified and folded into the dumbbell structure as previously described (Eustermann et al. 2011).

### Phosphorylation and ligation of nicked DNA ligand

The double-labeled DNA ligand was phosphorylated using the T4 Polynucleotide Kinase for 1 hour at 37°C, followed by heat inactivation. This 5’-phosphorylated version of the DNA ligand was used for all smFRET experiments.For ligation of the nick, the phosphorylated DNA was diluted to 250nM in T4 DNA Ligase Buffer and 2000 units of T4 DNA ligase were added in the reaction volume of 20μl. The reaction was incubated overnight at 16°C and then the ligase was heat inactivated.

### Protein expression and purification (PARP-1, F1F2, F2)

The codon optimized DNA plasmid for PARP-1 expression in E. coli was kindly provided by David Neuhaus. Expression and purification procedures were adopted with minor changes from protocols also provided by Sebastian Eustermann and David Neuhaus and protocols from Langelier 2011. Expression of PARP-1 was performed using E. coli BL21 (DE3) cells in Lysogeny Broth (LB) media supplemented with 50 μg/ml kanamycin and 10mM benzamide. All the colonies from a transformation plate were transferred to 60ml LBhiBenz media supplemented with 0.5% Glucose, 2mM MgSO4, 50 μg/ml Kanamycin to grow the starter culture to ≥2.0 OD600. 3x 1 l LBhiBenz were inoculated with 1:50 starter and allowed to grow to 0.8-1.2 OD600, 37oC, 200 rpm. Cell growth was arrested at 0.7 and incubated for 1 h at 4°C. Afterwards, expression was induced with 0.5mM IPTG and 0.15mM ZnSO4 and allowed to proceed overnight at 25°C, shaking at 200rpm. After centrifugation, the cell pellet was resuspended in 80 ml buffer containing 25mM HEPES-Na, pH 8, 0.5M NaCl, 0.5mM Tris(2-carboxyethyl)phosphine hydrochloride (TCEP), 1mM PMSF and 1 protease inhibitor tablet (EDTA free, Sigma) and sonicated for 3 minutes at 75% amplitude, 3/8 seconds on/off. After clearing the lysate, it was mixed with 8ml Ni-NTA beads (50% slurry, Qiagen) and incubated for 90 min on rollers at 4°C. The lysate mixture with Ni-NTA agarose was applied to a Bio-rad gravity column (2.5 × 10 cm). After the flow through was collected, the Ni-NTA agarose with bound PARP-1 was washed with 50ml LSW (low salt wash buffer containing 25mM HEPES-Na, pH 8, 0.5M NaCl, 20mM Imidazole, 1mM PMSF, 0.5mM TCEP) followed by 50 ml HSW (high salt wash buffer as LSW but with 1M NaCl) and again with 50 ml LSW. The elution was performed using 10 ml, 8ml, and 8 ml with 5 minutes incubation of elution buffer containing 25mM HEPES-Na, pH 8, 0.5M NaCl, 400mM Imidazole, 1mM PMSF, 0.5mM TCEP. After purification with Ni-NTA agarose the protein was diluted to 375mM NaCl and further purified on an ÄKTA purifier system (GE Healthcare Life Sciences) at 4 °C using 1ml HiTrap™ Heparin HP column (Cytiva). The protein was eluted in buffer containing 50 mM Tris, pH 7, 0.5mM TCEP and a salt gradient from 375mM to 1M NaCl. After this purification step the PARP-1 was subjected to buffer exchange (20mM Hepes pH 8.0, 200mM NaCl, and 0.1mM TCEP) with Zeba™ Spin Desalting Columns, 7K MWCO, 0.5 mL (Thermo Scientific).

Expression and purification of F2 was performed as described (Eustermann et al. 2011), with the following minor modifications: Expression was carried out in Rosetta2 cells (Novagen) and at 16°C; one tablet of SigmaFast-EDTA free protease inhibitors was used during expression, but no protease inhibitor was added during purification; ion exchange chromatography for initial protein purification was performed on a 1ml HiTrap SP FF (GE healthcare) and a 24ml Superdex 75 10 300 GL (GE healthcare) was used for size-exclusion chromatography. The respective protein sequences are given in Supp. Table S3.

### Single-molecule FRET measurements

#### Optical setup

SmFRET measurements were performed on a custom built confocal fluorescence setup using time-correlated single photon counting (TCSPC) and pulsed interleaved excitation combined with multiparameter fluorescence detection (PIE-MFD) (Kudryavtsev et al. 2012). The setup has recently been described in detail (Schwarz et al. 2018).

In brief, excitation lasers at 531 nm and 640 nm were pulsed with a repetition rate of 20 MHz and with a shift of about 20 ns between each other. The excitation light is focused into a sample droplet by a 1.2 NA water immersion objective. Excitation powers were 25 μW for the red laser and 95 μW for the green laser for measurements of the DNA labeled with 6-Tamra and Alexa647. For measurements of the alternative DNA labeled with Atto550 and Alexa647, intensities were 40 μW for the red and 95 μW for the green laser. Light emitted from single molecules diffusing through the confocal volume was collected by the same objective, focused onto a 75 μm pinhole and split into four detection channels according to polarization and spectral range. The light in each channel was focused onto a single-photon-counting avalanche photodiode and photon arrival times were recorded by TCSPC electronics.

#### Sample preparation

Prior to smFRET measurements with DNA+F1F2, DNA+F2 or DNA alone, the DNA was diluted in measurement buffer (12mM HEPES pH8.0, 60mM KCl, 3mM MgCl2, 50μM ZnSO4, 4% Glycerol, 0.5mM DTT) to a concentration of 5-30pM. Then, the respective PARP-1 fragment was added to a final concentration of 1μM in case of the DNA+F1F2 or 10μM in case of the DNA+F2 and then used for measurements directly. For smFRET measurements with DNA+PARP-1, 20pM of DNA in presence of 1μM protein in an optimized buffer (20mM HEPES pH 8.0, 200mM NaCl, 0.1mM TCEP) was used. The measurements with PARPi were performed in the same conditions as with PARP-1 but with addition of 200μM of either EB-47 or niraparib (MedChemExpress LLC, USA). PARPi stock solutions were prepared in DMSO such that the stock concentration allowed the DMSO percentage in smFRET measurements to be kept below 2%. All samples were kept on ice until start of the measurements, which took place at room temperature. A droplet of the sample was placed on a PEGylated coverslip (Marienfeld No. 1) and restrained using Roti Liquid Barrier Marker, colorless (Roth).

#### PEGylation of coverslips

Coverslips were first cleaned by cooking once in a 2% solution of Hellmanex (Hellma Analytics) and twice in water. They were then rinsed with water and dried under a nitrogen stream. The dry coverslips were treated in a PlasmaCleaner (Zepto, Diener electronic) for 10 minutes with oxygen plasma at 80% power. The clean coverslips were silanized by incubating in a solution of 2% (v/v) 3-aminopropyl-triethoxysilane (Sigma-Aldrich A3648) in acetone for 30 minutes. After washing and drying, the coverslips were PEGylated by incubation with a solution of mPEG-SVA (methoxy-poly(ethylene glycole)-succinimidyl valerate (Laysan Bio Inc. #MPEG-SVA-5000, 400 mg/ml in a 3:7 carbonate:bicarbonate buffer pH9.4) for 45 minutes. PEGylated coverslips were rinsed with water, dried under nitrogen stream and kept in a dry container until use (Kallis 2020).

### Data analysis

Data analysis was performed using a MATLAB based software package called PAM (PIE analysis with MATLAB) (Schrimpf et al. 2018). The newest version is available via a repository (https://gitlab.com/PAM-PIE/PAM).

#### Burst selection and quantification of burst-wise parameters

Single-molecule events (bursts) were identified in the photon time traces by applying an all-photon burst search (Nir et al. 2006), requiring at least 10 photons in 500 μs and more than 50 photons in total per burst. Files with more than 13 bursts per second were not considered for analysis to avoid frequent occurrence of multi-molecule events. All data sets were corrected for background, direct excitation (δ), crosstalk (α), γ (accounting for differences in quantum yield and detection efficiencies for the two dyes) and β (accounting for different absorption cross sections and laser intensities), following standard (Hellenkamp et al. 2018); the values determined are summarized in Supp. Table M1. Background count rates were determined from a measurement of pure buffer and ranged from 0.1 to 0.9 kHz, depending on polarization and color of the detection channel. Crosstalk was determined from the donor-only labeled species in the measurements. Direct excitation was quantified from an acceptor-only reference, comprising only the 5’ stem of the DNAAT550. γ and β were obtained from a fit of the stoichiometry vs. FRET efficiency histograms of high and low FRET species (DNA with and without protein, respectively), as described in (Lee et al. 2005; Kudryavtsev et al. 2012). For calculation of burst-wise anisotropies of the acceptor, further correction factors were taken into account, namely l1=0.03 and l2=0.090, accounting for depolarization introduced by the high-NA objective (Koshioka, Sasaki, and Masuhara 1995) and G=0.95, accounting for different detection efficiencies of the parallel and perpendicular polarization detection channels (Schaffer et al. 1999). Burst-wise lifetimes were calculated based on a maximum likelihood estimator (Maus et al. 2001). FRET events were isolated from the data using an upper threshold of 10 for the ALEX-2CDE filter described in (Tomov et al. 2012), which removes multi-molecule and blinking events. Remaining donor- or acceptor-only events were removed with a stoichiometry threshold (0.3<Sto<0.75). The presented FRET histograms contain only events with at least 100 photons and a corrected FRET efficiency between 0 and 1. The mean FRET efficiencies for each DNA conformation were quantified by fitting Gaussian functions to the peaks. The uncertainty for the experimental FRET efficiency was determined by error propagation of the uncertainties in the correction factors and background counts according to equation 1.

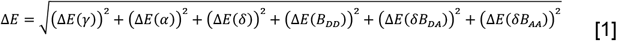

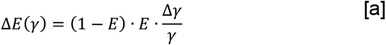

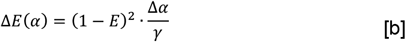

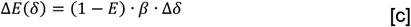

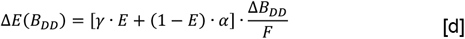

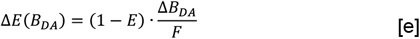

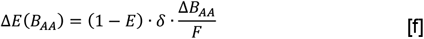

The relevant correction factors are summarized in Table M1. Following (Hellenkamp et al. 2018), we used relative errors of 10% for α, β and γ. For the mean photon number per burst F, we used 75 and an error in the background counts of 1 photon (ΔB_DD_, ΔB_DA_ and ΔB_AA_, for donor, FRET and acceptor channel, respectively). The resulting error ΔE depends on the measured FRET efficiency and ranged from 0.01 to 0.03 for the relevant range of FRET efficiencies, so we used ΔE=0.03 as upper estimate.

**Table M1:**
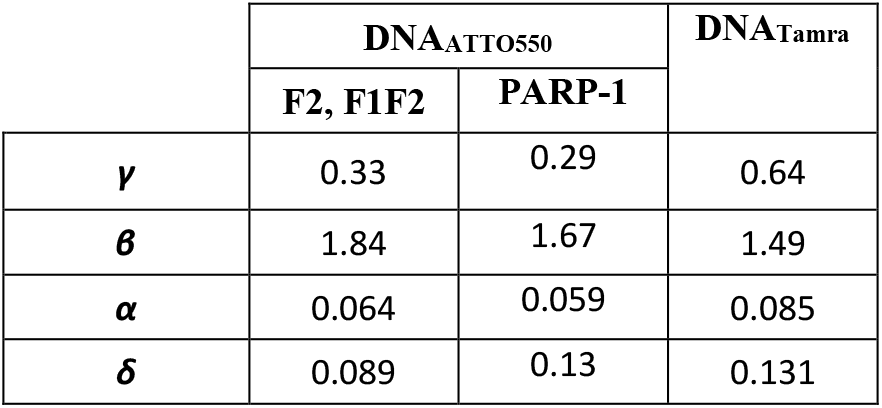
Correction factors applied for the two DNA construct.

### Generation of model ensembles

Models were calculated with the program XPLOR-NIH (Schwieters et al. 2003) using a slightly adapted version of the procedure previously described in detail in (Eustermann et al. 2015) Supplemental Experimental Procedures. First, template structures were calculated for the DNA and protein components. The protein template structures were calculated starting from the deposited co-ordinates of PARP-1 F1 and F2 domains in complex with DNA blunt ends; chains B, I and J from pdb 3ODA were used for the complex of F1, and chains B, E and F from pdb 3ODC were used for the complex of F2. These co-ordinates were adapted to the XPLOR-NIH force field by addition of hydrogen atoms followed by energy minimization while holding all the backbone peptide groups fixed. A starting DNA dumbbell template structure was generated by taking the lowest energy structure from an ensemble of 50 simulated annealing structures calculated using a combination of backbone intra-strand and inter-strand distance restraints, base pairing hydrogen-bond distance restraints and dihedral angle restraints for ideal B-form geometry in the stems (all measured from ideal structures generated using the webserver at http://structure.usc.edu/make-na) as well as weak base-pair planarity restraints, and using distance and dihedral angle restraints corresponding to measurements from RNA tetraloop structures 1MSY and 1RNG for the first and second tetraloops respectively (as in the previous work, the ring of the second T in the second tetraloop was modeled in the anti-conformation). The resulting DNA dumbbell structure served as the template for ensembles containing DNA only. For ensembles that included, in addition to the DNA, either F2 alone (bound to the DNA 3’ stem) or both F1 and F2 (bound to the 5’ and 3’ DNA stems respectively), those DNA stems that were to be bound to protein were adapted to mimic very closely the conformations found in the corresponding DNA stems in 3ODA (for the 5’ stem) and 3ODC (for the 3’ stem). This was achieved by simulated annealing calculations during which the ideal B-form backbone-backbone distance restraints and dihedral angle restraints in the 7-basepair regions bound by protein were replaced by corresponding restraints using values measured in 3ODA or 3ODC; during this process, the remaining portions of each dumbbell were restrained using the same ideal B-form restraints within the stems and tetraloop restraints as described above. The only adaptations required for the present calculations relative to those described in (Eustermann et al. 2015) comprised the use of these B-form restraints in stem regions between the protein binding site(s) and the tetraloops, required here because the stems being modeled were longer, as well as the absence of the single nucleotide linker that had been present between the 3’ and 5’ stems in the earlier calculations.

Ensembles were generated using these template structures by first randomizing the five rotatable dihedral angles of the 3’ stem to 5’ stem linker in the DNA dumbbell, then adding the template protein structures for those stems that were to be protein-bound, using for fitting the DNA backbone atoms in the 7-basepair region of each protein domain’s DNA-binding footprint that were in common (these atoms had also been taken from 3ODC and 3ODA during generation of the protein templates); in this way, each protein domain was carried accurately into the same spatial relationship to the appropriate stem of the dumbbell as it had with blunt-ended DNA in complex 3ODA (for F1) or 3ODC (for F2). Structures were then subjected to long simulated annealing calculations with small step sizes, during which the conformation of the protein domains, the DNA stems and tetraloops, but crucially not the DNA linker, were restrained using non-crystallographic symmetry (NCS) restraints relative to rigidly constrained copies of the starting template structures; this procedure is described in detail in (Eustermann et al. 2015) Supplemental Experimental Procedures. In the case of the F1F2-DNA ensemble, the exact same NMR-derived restraints were applied during this annealing process as had been used in (Eustermann et al. 2015), that is a combination of NOE-derived interdomain and intermolecular distance restraints and residual dipolar coupling-derived orientational restraints as measured for the experimentally determined complex of F1F2 with a 45-nucleotide dumbbell. In the cases of the F2-DNA ensemble and the DNA-only ensembles, no NMR-derived restraints were applied during the annealing calculations. For the F1F2-DNA ensemble, 500 structures were calculated, from which the best 49 were selected according to three simultaneously applied selection criteria based on XPLOR-NIH energy terms (these criteria were E(total) < 6000 kcal.mol-1, E(NOE) < 30 kcal.mol-1 and E(tensor) < 1400 kcal.mol-1). For the F2-DNA and DNA-only ensembles, in order to sample conformation space adequately it was found necessary to select the 1000 structures with lowest XPLOR-NIH E(total) values from a total of 5000 calculated for each ensemble.

Finally, in the cases of the F1F2-DNA and F2-DNA ensembles, the additional atoms of the flexible N- and C-terminal protein tails and, in the case of the F1F2-DNA ensemble only, also the F1F2 linker (none of which were present in the protein template structures), were added using a further simulated annealing protocol, during which the ordered portion of the structure (the whole DNA except for the 5’ phosphate group, as well as residues 6-91 of F1 and 109-201 of F2) was held rigid.

### Structural analysis

#### FRET efficiency simulation

Our approach uses an ensemble of possible conformations of the macromolecular complex. For every structure, we compute the AVs of donor and acceptor by using the structure analysis tool FastNPS (Eilert et al. 2017). As a user input, the algorithm needs the attachment point and the geometry of both the fluorophore and its linker, as well as the pdb-file of the structural ensemble. The dye and linker parameters were determined from the chemical structure using Chem3D 16.0 (PerkinElmer) and are summarized in Supp. Table S4. With this information, FastNPS computes the volume which is accessible to the dye. These volumes are saved and used for the stochastic simulation of the FRET process. As described in (Eilert et al. 2018), it is assumed that the dyes can translate freely within the AVs. For rotation, we propose that the dyes diffuse in a spherical cone with random axis obtained from the AV by a principle component analysis. As the diffusion coefficient of a dye relative to the molecule cannot be extracted from the smFRET measurements, a recently reported value of 10 Å^2^/ns is assumed for all dyes (Peulen, Opanasyuk, and Seidel 2017). The semi-angle of the cone and the rotational diffusion coefficient are determined from time-resolved anisotropy data (Supp. Table S5). For every conformation, we simulate 50000 trajectories, i.e. the translational and rotational diffusion of donor and acceptor, and obtain an average smFRET efficiency. Using 50000 trajectories per structure reduced the standard deviation of 100 independent simulations for the same model to 0.2% (as compared to 0.5% for 10000 or 1.7% for 1000 simulated excitations), so that the statistical error is small compared to the overall uncertainty of the simulations. The parameters used in the simulations are given in Supp. Table S4 and Supp. Table S5. Details about their determination are given in the section “Data analysis” and “Determination of the isotropic Förster radius”.

#### Selection and analysis of structures

We select conformations whose simulated efficiencies are within an error of ΔE=0.05 compared to the experimental result. This accounts for both the error in the isotropic Förster Radius (ΔRiso=2Å translates to ΔE=0.04 in the simulations, see also section “Determination of the isotropic Förster radius”) and the error of the measurement derived from the uncertainties in the correction factors (ΔE=0.03, see section “Burst selection and quantification of burst-wise parameters”). Every linear dsDNA molecule has a unique axis. We are interested in the angle between the axes of the 5’- and 3’-stems. As a convention, the direction of their axes is defined from the nick to the respective loop. We define the axis of each DNA stem as the longest axis determined from a principal component analysis, considering all atoms belonging to the respective stem, but omitting the four bases of the tetraloop and the base pair next to the loop. All structures are aligned with respect to the 3’-stem, such that we can represent every selected conformation as a unit vector/orientation (representing the 5’-stem) in a spherical coordinate system. This allows us to compute the angle between the axes of the two stems for every conformation. We obtain histograms of these angles where we re-weight the relative frequency in the equally spaced bins according to the differing surface of the ring on the unit sphere. For the computation of mean angles and standard deviations we considered two axially symmetric scenarios: the orientations lie in a spherical cone or in an open spherical sector (the symmetric difference of two cones with same mean axis). In the view along the mean axis, they represent a circle and an annulus, respectively (Supplementary Fig. S12). In case of a spherical cone, we have a single mean axis which is computed by the normalized sum over all axes. The standard deviation is retrieved from the corrected histogram. If the orientations lie within an open spherical sector, we do not have one unique mean orientation, but it is axially degenerate, i.e. it is on a lateral surface of a cone. However, the mean polar angle between the stems is still uniquely defined, if we assume that the open spherical sector is directed along the 3’-axis. Thus, both the mean angle and the standard deviation are computed by the corrected histogram.

#### Visualization of structures

Pictures of molecular structures were generated using the UCSF Chimera package (Pettersen et al. 2004). Chimera is developed by the Resource for Biocomputing, Visualization, and Informatics at the University of California, San Francisco (supported by NIGMS P41-GM103311).

### Filtered FCS

Filtered FCS (fFCS) was employed to disentangle the mixture of species and to filter out the separate species using their difference in FRET efficiency (Felekyan et al. 2012). Thus to separate the interconverting states in DNA+F2 and DNA+F1F2 datasets, burstwise fFCS with a time window of 50 ms around the edges of each burst was performed using the MATLAB package PAM (Schrimpf et al. 2018) as described in (Barth et al. 2018). The bursts corresponding to separate species (states) were selected by setting the respective FRET efficiency thresholds (species 1: 0.25<E<0:35, species 2: 0.77<E<0.87 for DNA+F2; species 1: 0.34<E<0:42, species 2: 0.9<E<0.98 for DNA+F1F2, Fig. 5). For every species, TCSPC patterns in donor and FRET channels were plotted from which the stacked photon histograms - fFCS filters were generated. The filters of each species were used to calculate the auto- and cross-correlation functions (Felekyan et al. 2012). For each dataset (DNA+F2 and DNA+F1F2) four correlation functions were calculated, namely species 1 x species 1, species 1 x species 2, species 2 x species 1, species 2 x species 2. The correlation functions were fit with globally with a diffusion (G_diff_) model containing two kinetic terms (τ_Ri_):

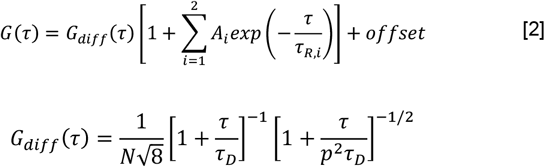

## End Matter

### Author Contributions and Notes

J.M. and S.E. designed the study. A.S. and E.K. performed the experiments and analysed the data. T.E developed the angle quantification analysis. C.R. provided technical support for the experimental setup for the smFRET analysis. D.N generated all the structural ensembles and provided the plasmid DNA for PARP-1. A.S. and S.E. expressed and purified proteins. E.K, A.S, J.M, D.N and S.E. wrote the manuscript.

## Acknowledgments

We are greatful for the support from Nadine Jakobi performing the labeling of the DNA donor strand, Mara Guriento who performed F2 purification and Laura Easton who provided the protocol for PARP-1 purification in *e*.*Coli*.

## Funding

This work was supported by the Deutsche Forschungsgemeinschaft through the CRC 1279 (JM). DN was supported by the Medical Research Council [U105178934]. EK acknowledges support by the Carl-Zeiss Stiftung.

## Data Availability

All data is available online through the Dryad Digital Repository (https://datadryad.org/stash/share/Bb3aS7D8DXMfGRX9gSCi44DzH-Bkix_0ZwsafCp5yuY). The MATLAB algorithm for DNA kink angle from smFRET analysis is available at https://github.com/Michaelislab/-DNA-kink-angle-from-smFRET-analysis.

## Competing interests

The authors declare that there are no competing interests.

## Notes

https://datadryad.org/stash/share/Bb3aS7D8DXMfGRX9gSCi44DzH-Bkix_0ZwsafCp5yuY

https://github.com/Michaelislab/-DNA-kink-angle-from-smFRET-analysis

### Competing Interest Statement

The authors have declared no competing interest.

