## Supplementary Information for "Structural dynamics of DNA strand break sensing by PARP-1 at a single-molecule level"

### Supplementary Methods

#### Gel electrophoresis

The ligation efficiency was assessed via gel electrophoresis. Phosphorylated DNA, ligated DNA and the control lacking the ligase were loaded on a 15% PAA 1xTBE 7M urea gel (16x20cm). The gel was prerun for 1.5 hours at 300 V and run for 8.5 hours. The gel was imaged on a ChemiDoc MP imaging system (BioRad)

#### Determination of the isotropic Förster radius

In order to quantify the isotropic Förster radius, several parameters had to be measured for the dyes in context of the DNA construct. The quantum yield of the donor attached to the DNA in absence of the acceptor was determined by product of the lifetime (see section lifetime and time-resolved anisotropy fits) and the radiative rate constant. The lifetime of the donor in absence of the acceptor was obtained from the subensemble analysis of the donor-only population of the smFRET measurement of donor only strand for $DNA_{Atto550}$ (methods) and from smFRET measurement of $DNA_{Tamra}$. For the radiative rate constant of Atto550 the value given by the manufacturer was used. As for 6-Tamra, the radiative rate constant was quantified by measuring the lifetime and quantum yield of unconjugated 6-Tamra. To calculate the spectral overlap *J* for $DNA_{Atto550}$ the emission of the donor strand was recorded. For the alternative DNA construct, $DNA_{Tamra}$, to measure the emission spectrum of the donor in absence of FRET, a reference sample (double stranded DNA, internally labeled with 6-Tamra and diluted in the buffer used for smFRET measurements) had to be used. Both donor emission spectra were recorded on a SPEX Fluorolog II (Horiba) (spectral bandwidth of 4.25nm for the excitation and 2.13nm for the emission) under magic angle conditions. For measuring the absorption spectrum of the acceptor, a DNA strand comprising only the acceptor labeled 3' stem of the $DNA_{AT550}$ (see section on synthesis of the DNA ligand) was diluted in the buffer also used for smFRET measurements. The spectrum was taken on a Cary 50 Bio spectrophotometer (Varian). Both spectra are needed to determine the overlap integral. Additionally, the absorption coefficient of Alexa647 at the absorption maximum (651nm determined from the measured spectrum) is needed and given as 270000 $M^{-1}cm^{-1}$ by the manufacturer (ThermoFisher Scientific). A value of 1.35 was assumed for the refractive index of the medium between the two dyes. PhotochemCAD 3 (Taniguchi, Du, and Lindsey 2018) was used to calculate the isotropic Förster distance $R_{iso}$=70Å for $DNA_{Atto}$ $R_{iso}$=68Å for $DNA_{Tamra}$ (Kallis 2020) from the determined parameters.

The uncertainty for the isotropic Förster radius $\Delta R_{iso}$ was determined according to equation 1 by error propagation of errors in the refractive index $n$, the donor quantum yield ϕ and the overlap integral $J$.

$$\Delta R_{iso} = \sqrt{\left(\Delta R_{iso}(n)\right)^2 + \left(\Delta R_{iso}(\Phi)\right)^2 + \left(\Delta R_{iso}(J)\right)^2} \qquad (1)$$

Equations for $\Delta R_{iso}(n)$, $\Delta R_{iso}(n)$ and $\Delta R_{iso}(n)$ are given in (Hellenkamp et al. 2018). Due to the position of our labels on the DNA, we expect that the medium between them will be composed almost exclusively of buffer. We thus assume a refractive index of $n$=1.35±0.02, which leads to $\Delta R_{iso}(n)$=0.01*$R_{iso}$. We further assumed an error of 4% for the donor quantum yield based on the uncertainties in the donor lifetime and radiative rate constant, leading to $\Delta R_{iso}(\phi)$=0.01*$R_{iso}$. The value of $\Delta R_{iso}(J)$=0.025%*$R_{iso}$ was adopted from (Hellenkamp et al. 2018). These uncertainties lead to $\Delta R_{iso}$=0.03*$R_{iso}$=2Å for the construct $DNA_{Tamra}$.

<u>Data for simulations: lifetime and time-resolved anisotropy</u>

Data for the donor in absence of the acceptor was obtained from the donor-only population of the smFRET measurements. All donor photons after donor excitation (531nm) of the donor-only population (uncorrected stoichiometry >0.9 and more than 100 photons) were combined in one TCSPC histogram. Data for the acceptor can be gained either from the acceptor-only population (uncorrected stoichiometry <0.25 and more than 70 photons) or from the FRET population (as defined above), because the acceptor photons after acceptor excitation (640nm) are not affected by the FRET process. Results for both species were generally in good agreement, so a mean value was taken.

Fluorescent lifetime, rotational correlation time and residual anisotropy were determined by iterative re-convolution fitting of the parallel and perpendicular decays (Supplementary Fig. S11). Concentrated solutions of crystal violet or malachite green (two fluorescent dyes with lifetimes in the picosecond range) were used to record the instrument response function (IRF) of the green and red channels, respectively.

In a first step, the fluorescence lifetime was determined from the combined TCSPC histogram $D(t)$, which adds the data from parallel ($D_{\parallel}$) and perpendicular ($D_{\perp}$) detection channels according to

$$D(t) = (1 - 3l_2) \cdot G \cdot D_{\parallel}(t) + (2 - 3l_1) \cdot D_{\perp}(t) \qquad (2)$$

with $G$, $l_1$ and $l_2$ being the correction factors described above. For Atto550 a monoexponential and a biexponential model function were needed to describe the acceptor Alexa647. Iterative re-convolution of the model decay with the recorded IRF was used to fit the TCSPC histogram $D(t)$, additionally considering a constant background contribution. For the simulations, the amplitude weighted average lifetime of the two donor components was used. The value for the donor lifetime given in Supp. Table S5 represents the average of the fit results from DNA, DNA+F2 and DNA+F1F2, because they were in agreement.

In the second step, the anisotropy information was analyzed in a global fit of the TCSPC data from the parallel ($D_{\parallel}$) and perpendicular ($D_{\perp}$) detection channels by iterative re-convolution. The model functions were

$$M_{\parallel}(t) = \frac{1}{G} \cdot I(t) \cdot [1 + (2 - 3l_1) \cdot r(t)] \tag{3a}$$

$$M_{\perp}(t) = I(t) \cdot [1 - (1 - 3l_2) \cdot r(t)] \tag{3b}$$

with $I(t)$ being the biexponential fluorescence intensity decay and $r(t)$ being the model function for the anisotropy decay, given by one of the following functions, depending on the complexity of the data.

$$r(t) = (r_0 - r_\infty) \cdot e^{-\frac{t}{\rho}} + r_\infty \tag{4a}$$

$$r_{tumb}(t) = \left[(r_0 - r_\infty) \cdot e^{-\frac{t}{\rho}} + r_\infty\right] \cdot e^{-\frac{t}{\rho_{tumb}}} \tag{4b}$$

With the fundamental anisotropy $r_0$, the residual anisotropy $r_\infty$, the rotational correlation time of the dye $\rho$ and the tumbling time of the DNA $\rho_{tumb}$. In order to reduce the number of free parameters, the lifetimes and their fractions were fixed to the results from the lifetime fit. A constant background contribution was also assumed. The simple anisotropy model was used for the donor data of all datasets. For all acceptor data, the anisotropy model with two lifetimes and two rotational correlation times was used.

As in the case of the donor lifetime, the values given in Supp. Table S5 for the rotational correlation times of donor or acceptor represent the average of the fitting results from DNA, DNA+F2 and DNA+F1F2, because their difference was close to the expected uncertainty. The value for the residual anisotropy of the donor was taken from the fit of DNA+F1F2 (Supplementary Fig. S11), because for this dataset, the DNA tumbling time was sufficiently long to play no role. Thus, the residual anisotropy should be most accurate if determined from this dataset, where a simple model is sufficient. The value for the residual anisotropy of the acceptor was taken as the average of the fit results from DNA, DNA+F2 and DNA+F1F2. In the presence of protein, the situation becomes more complicated than our simulation can describe, with two acceptor states due to PIFE (Stennett et al. 2015). Even so, on a long timescale, the anisotropy is expected to decay to the lower value of the unrestricted state, which is likely to be dominant in the absence of protein.

Static and dynamic FRET lines

In order to test for conformational dynamics, static and dynamic FRET lines were simulated (Kalinin et al. 2010) and overlayed with the plot of FRET efficiency vs. burst-wise donor lifetime (Supplementary Fig. S5a). The required donor lifetime was obtained from the donor-only population of the measurement, the Förster radius for $DNA_{AT550}$ was 70 Å and the apparent linker length of 7 Å was determined from the measurement of ligated DNA by fitting the fluorescence decay by a Gaussian distance distribution. For the dynamic FRET line, interconversion was assumed between the state of free DNA and the most kinked state observed in the measurement of DNA+F1F2.

Burst variance analysis

Burst variance analysis (BVA) tests for dynamics by comparing the shot-noise-limited standard deviation of the mean proximity ratio within a burst to the observed standard deviation (Torella et al. 2011). Here, the $E_{PR}$ fluctuations are assessed via splitting the burst into chunks of 10 photons and then calculating the $E_{PR}$ for each chunk and the standard deviation over all chunks (burst-wise $\sigma$ of $E_{PR}$). A binned standard deviation is calculated over all chunks in all bursts in an $E_{PR}$-bin (bin size of $E_{PR}$ = 0.05, >50 bursts per bin). The $\sigma$ of $E_{PR}$ expected from shot-noise for each chunk is calculated and then compared to the binned $\sigma$ of $E_{PR}$. The bursts with $\sigma$ of $E_{PR}$ values close to the expected $\sigma$ (those that lie on the black line) indicate static FRET events, whereas those which are above the black line indicate dynamic FRET events (Supplementary Fig. S5b).

Time window analysis

Dynamic interconversion can be overseen if it takes place on the faster timescale than the observation time window (Gopich and Szabo 2007). To observe this dynamics, the time binning of the data was varied from 0.25 to 10 ms. If dynamics is present, the shape of E histograms changes upon increase of the time window length. Therefore, to provide additional evidence for conformational dynamics, FRET efficiency was plotted at 0.25, 0.5, 1, 2, 5, and 10 ms observation time windows (Supplementary Fig. 5c). A threshold of 50 photons was set to exclude the time windows with lower number of photons.

FRET-FCS of the acceptor only species

FRET-FCS (Schwille, Meyer-Almes, and Rigler 1997; Torres and Levitus 2007)analysis was employed to observe the diffusion time $\tau_D$ in low FRET populations of DNA+PARP-1 and DNA+PARP-1+niraparib (Supplementary Fig. S7). For this the red parallel and red perpendicular channels ($AA_{par}$, $AA_{per}$) of the respective low FRET populations (0<E<0.59 for DNA+PARP-1 and 0<E<0.55 for DNA+PARP-1+niraparib, 0.3<S<0.8 for both) were correlated.

The obtained autocorrelation functions were fit with a one component FCS diffusion model with an additional term, accounting for acceptor photophysics $\tau_{phot}$ with fraction $A_{phot}$:

$$G(\tau) = \frac{1}{N\sqrt{8}}\left[1+\frac{\tau}{\tau_D}\right]^{-1}\left[1+\frac{\tau}{p^2\tau_D}\right]^{-1/2}\left[1+\frac{A_{phot}}{1-A_{phot}}\ exp\left(-\frac{\tau}{\tau_{phot}}\right)\right]$$

where, N is the average number of the molecules in the effective observation volume, $\tau_D$ is the diffusion time (or dwell time in the observation volume, depends on radius $\omega_0$ of the observation volume, $\tau_D = \omega_0^2/4D$, where D is the diffusion coefficient, Ries and Schwille 2012), $p$ is the ratio of the lateral and axial dimensions of the observation volume.

Additionally, FRET-FCS analysis was employed to observe the effect of acceptor photophysics upon PARP-1 binding to DNA (Supplementary Fig. S6). To this end, the red parallel and red perpendicular channels of the acceptor only species ($AA_{par}$, $AA_{per}$), selected using a stoichiometry cut 0<S<0.23, from smFRET measurement) for ligated DNA, DNA, DNA+F2, DNA+F1F2 and DNA+PARP-1 were correlated. The obtained autocorrelation functions were fit with the same model described above model. The returned , $\tau_D$ and $\tau_{phot}$were fixed when fitting the fFCS curves (Methods).

**Supplementary figures**

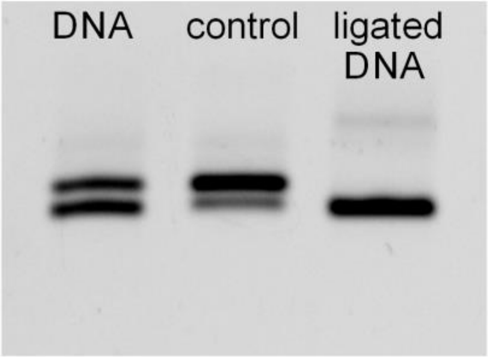


**Supplementary Figure S1:** Ligation efficiency of the nick was tested using gel electrophoresis

Gel electrophoresis of nicked and ligated DNA. DNA: phosphorylated nicked DNA before ligation. Ligated DNA: the same DNA after ligation with T4 DNA ligase. Control: the same DNA, treated as for the ligated sample except that water was added instead of ligase. The picture shows a FRET image, i.e. only double-labeled bands are visible. The two bands in the cases of the DNA and control samples are presumably due to the phosphorylated and unphosphorylated DNA fractions, the difference in the relative intensity between DNA and control could be explained by an additional phosphorylation step in the control sample (Methods). The gel demonstrates that ligation was very efficient.

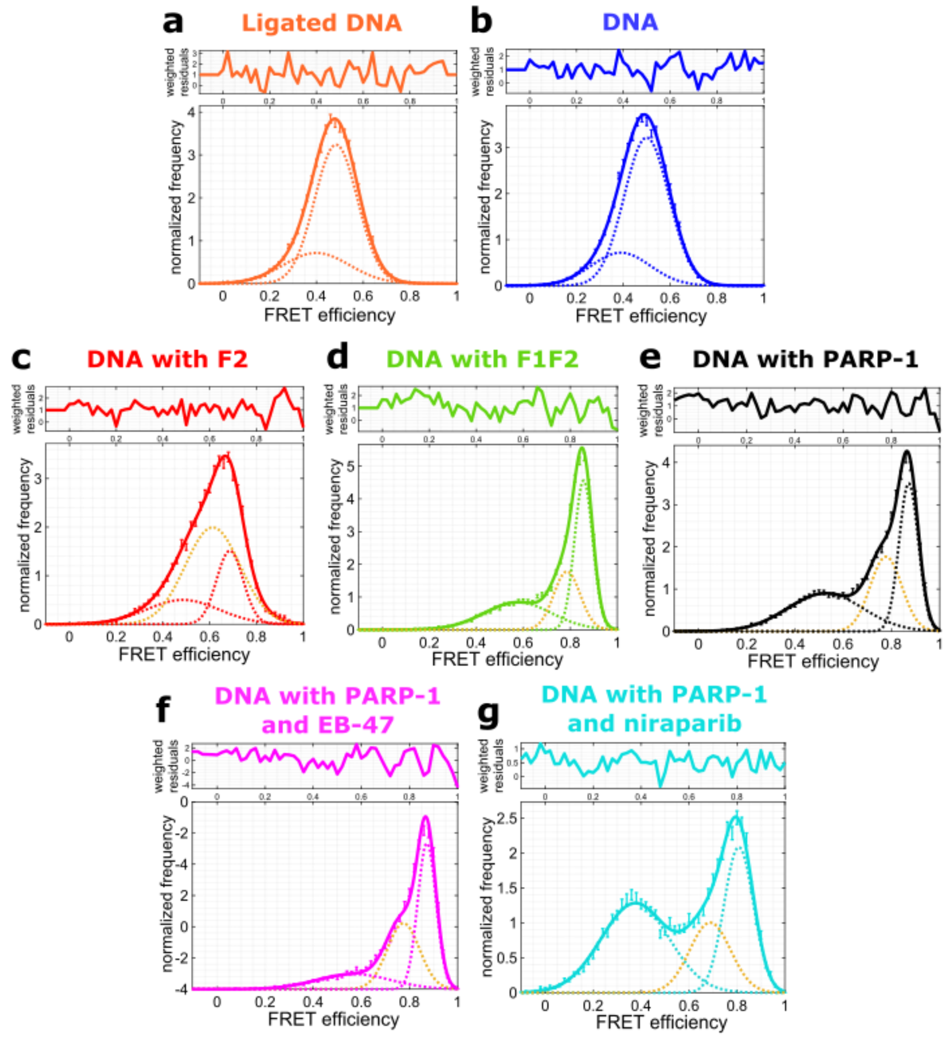


**Supplementary Figure S2: Quantification of FRET efficiencies.**

Gaussian fits of the smFRET efficiency histograms shown in Figs. 1, 2 and 6. **a:** Ligated DNA alone. **b:** DNA alone. **c:** DNA in presence of 10µM F2. **d:** DNA in presence of 1µM F1F2. **e:** DNA in presence of 1µM full-length PARP-1. **f:** DNA in presence of 1µM PARP-1 and 200µM EB-47. **g:** DNA in presence of 1µM PARP-1 and 200µM niraparib. The data was fitted with a two Gaussians (a and b) or with the sum of three Gaussians (b-g), where in each case the third Gaussian (yellow) accounts for the dynamic population. All parameters were freely adjustable during fitting. Fit results are summarized in Supp. Table 1.

**a**

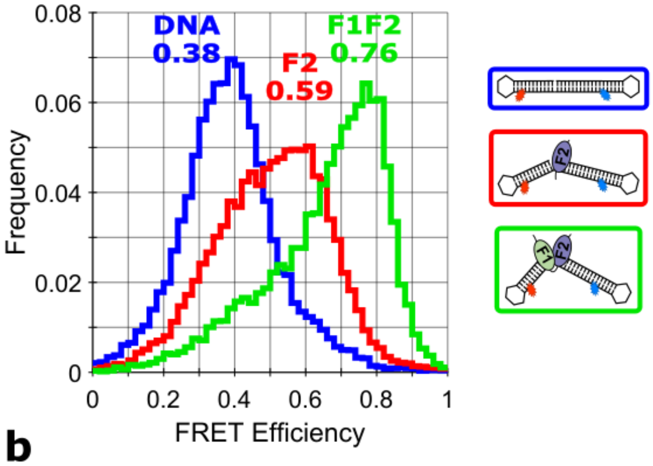


**b**

| | $A_1$ | $E_1$ | $\sigma_1$ | $A_2$ | $E_2$ | $\sigma_2$ | $R^2$ |
|---|---|---|---|---|---|---|---|
| **DNA** | 1.00 | 0.38 | 0.11 | | | | 0.986 |
| **lig. DNA** | 1.00 | 0.38 | 0.12 | | | | 0.988 |
| **F2** | 0.40 | 0.38 (fix) | 0.11 | 0.60 | 0.59 | 0.10 | 0.995 |
| **F1F2** | 0.51 | 0.57 | 0.17 | 0.49 | 0.76 | 0.08 | 0.985 |

**Supplementary Figure S3: DNA conformation during nick recognition probed by smFRET with an alternative DNA construct with TAMRA as donor ($DNA_{Tamra}$).**

In order to check the influence of dye molecules on the obtained results, experiments were repeated using an alternative DNA construct with TAMRA as a donor (Kallis 2020). **a:** smFRET efficiency histogram of nicked DNA (blue) and nicked DNA in presence of either 10µM F2 (red) or 1µM F1F2 (green). The indicated peak FRET efficiencies were obtained by Gaussian fitting. **b:** Results of Gaussian fitting of smFRET efficiency histograms where *A*: area fraction, *E*: mean FRET efficiency, *σ*: standard deviation of the Gaussian. FRET efficiencies of the species shown in A are highlighted. While the specific FRET efficiencies are slightly different the general effect observed is comparable to that of the original DNA construct.

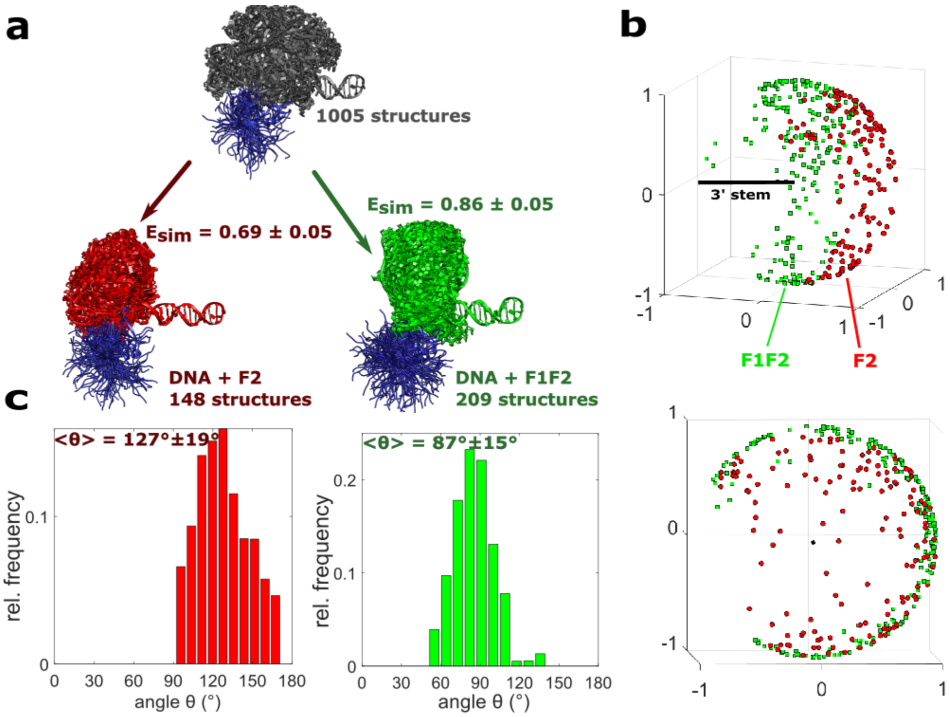


**Supplementary Figure S4: Selection and analysis of structures in accordance with experiment.**

**a:** Structures in agreement with the experimental FRET efficiencies (see Fig. 2) were selected from the ensemble of DNA+F2 (1005 structures, 100 of which are shown here in grey) as follows: $E$=0.69 for DNA+F2 (148 selected structures shown in red), $E$=0.86 for DNA+F1F2 (209 selected structures shown in green). An uncertainty of $\Delta E = \pm 0.05$ was assumed in all cases. **b:** Combined representation of the selected structures from DNA+F2 ensemble (1005 structures) shown in A. All structures are aligned with respect to the 3' stem, the axis of which is represented here by a black line. The colored shapes indicate the direction of the 5' axis for each selected structure: red circles for DNA+F2; green squares for DNA+F1F2. Two different views of the 3D plot are shown, a side view (top) and a view along the 3' axis (bottom). **c:** Kinking angle distributions for the selected structures in a: DNA+F2 (red, left hand histogram), DNA+F1F2 (green, right hand histogram). The mean and standard deviation of the angles are indicated for each histogram.

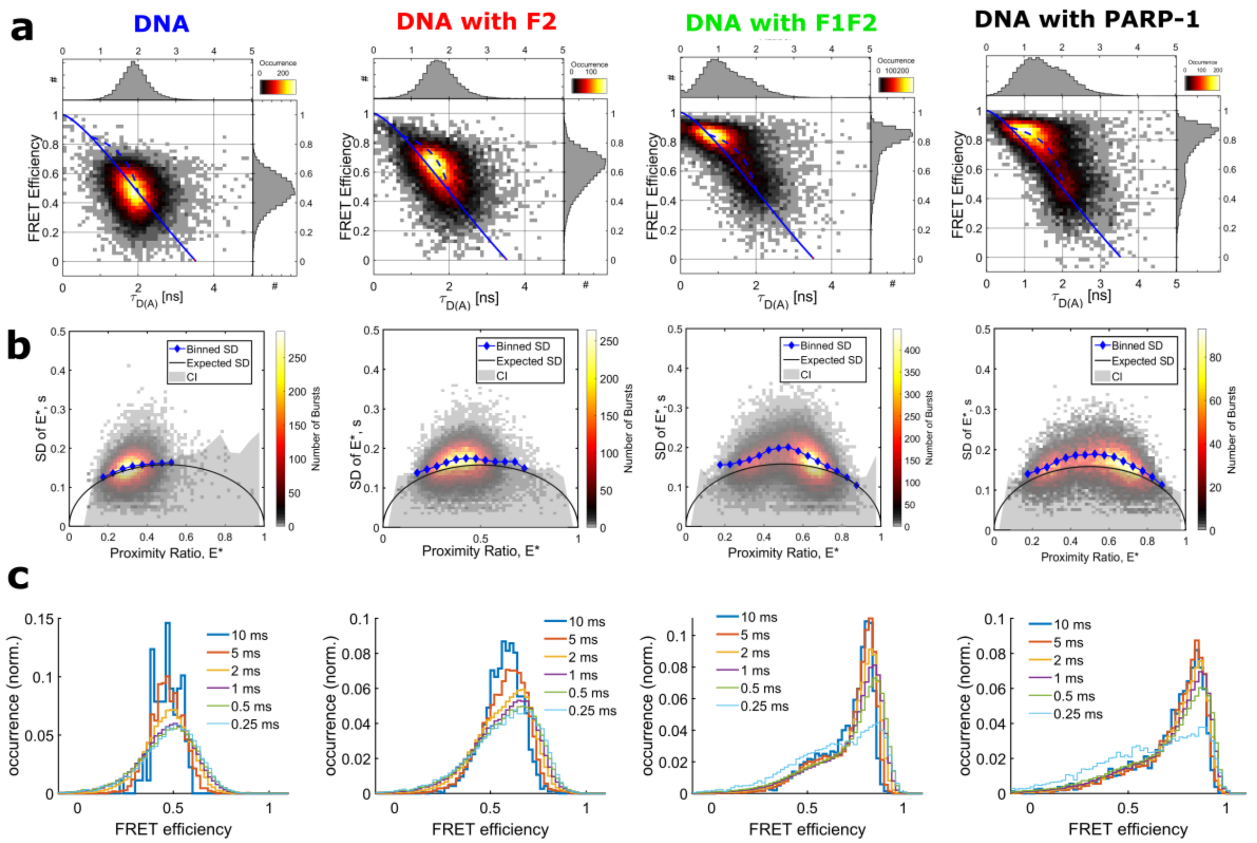


**Supplementary Figure S5**: **Qualitative analysis of dynamics for DNA, DNA+F2, DNA+F1F2 and DNA+PARP-1 datasets**

**a:** E vs. TauD plots with static (solid) and dynamic (dashed) lines indicate dynamics for DNA with F2, DNA with F1F2 and DNA with PARP-1 where the respective population deviates from the static FRET line. **b:** Burst variance analysis (BVA) results show the dynamics in DNA with F2, DNA with F1F2 and DNA with PARP-1 where the binned standard deviation (blue diamonds) is above the expected standard deviation (black solid line). **c:** Time window analysis results show the dynamics in DNA with F2, DNA+F1F2 and DNA with PARP-1 where the FRET efficiency histograms change their shape upon increase of the time window. All three analysis schemes (Supplementary methods) indicate the presence of the dynamic interconversion in the cases of DNA bound by PARP-1 or its fragments, whereas in contrast for the DNA alone no dynamics is observed.

**a**

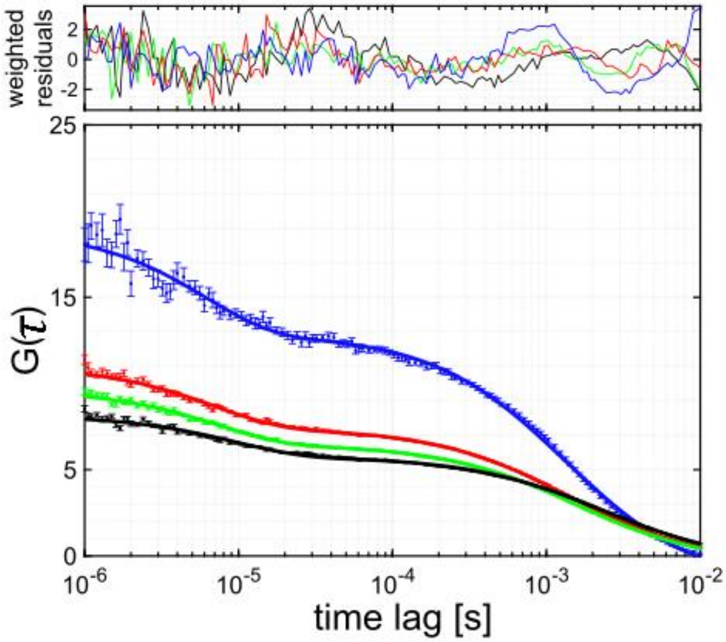


**b**

| Data Set | N | $\tau_D$ [µs] | $\tau_{phot}$ [µs] | $A_{phot}$ | $y_0$ | $\chi^2$ |
|---|---|---|---|---|---|---|
| DNA | 0.024 | 1341 | 6.08 | 0.43 | -1.52 | 1.60 |
| DNA with F2 | 0.052 | 1411 | 7.01 | 0.47 | -0.30 | 1.07 |
| DNA with F1F2 | 0.046 | 1587 | 7.38 | 0.47 | -0.37 | 0.89 |
| DNA with PARP-1 | 0.058 | 2342 | 9.58 | 0.40 | -0.29 | 1.86 |

**Supplementary Figure S6: Investigation of the acceptor blinking time $\tau_{phot}$.**

**a:** The respective fitted autocorrelation functions of acceptor only species in the red channel: blue – DNA, red – DNA+F2, green – DNA+F1F2 and black – PARP-1 which was correlated additionally for better comparison. The upper panel shows the weighted residuals for each correlation function. **b:** Corresponding fit results for autocorrelation functions of the acceptor only species in the red channel shown in b. The diffusion time $\tau_D$ and the acceptor blinking component $\tau_{phot}$ are used in the fFCS (Methods) as fixed parameters. An increased $\tau_{phot}$ is observed for DNA bound by protein datasets . The increase of $\tau_{phot}$ here occurs most probably due to protein induced fluorescence enhancement (PIFE) effect (Stennett et al. 2015) as our acceptor dye is attached to 5' stem, exactly where the PARP-1 or its fragments bind.

**a**

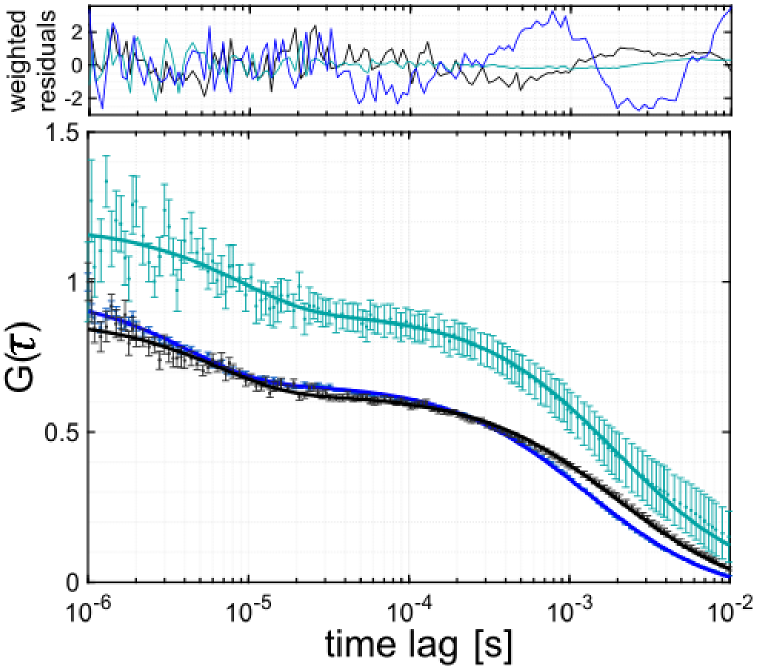


**b**

| Data Set | N | $\tau_D$ [µs] | $\tau_{phot}$ [µs] | $A_{phot}$ | $y_0$ | $\chi^2$ |
|---|---|---|---|---|---|---|
| **DNA** | 0.027 | 1303 | 4.30 | 0.30 | -0.96 | 2.50 |
| **DNA with PARP-1** | 0.039 | 1990 | 6.48 | 0.27 | -0.72 | 0.78 |
| **DNA with PARP-1 and Nirapairb** | 0.028 | 1976 | 8.88 | 0.24 | -013 | 0.34 |

**Supplementary Figure S7: Investigation of diffusion time $\tau_D$ for low FRET species of DNA+PARP-1 and DNA+PARP-1+Niraparib.**

**a:** Fitted autocorrelation functions of FRET sub-species: blue – DNA, grey – low FRET population of the DNA+PARP-1 (0 – 0.59), turquoise – low FRET population of the DNA+PARP-1+niraparib (0 – 0.55). The upper panel shows the weighted residuals for each correlation function. For the fits data from the acceptor fluorescence after acceptor excitation was used. **b:** Corresponding fit results for the respective autocorrelation functions: N is the average number of molecules in observation volume, $\tau_D$ is the diffusion time, $\tau_{phot}$ is a time scale characterizing acceptor blinking (compare to Supplementary Fig. S6), $A_{phot}$ is the fraction of the blinking and $y_0$ is the offset (Methods). An increased $\tau_{phot}$ is observed for DNA bound by protein datasets .

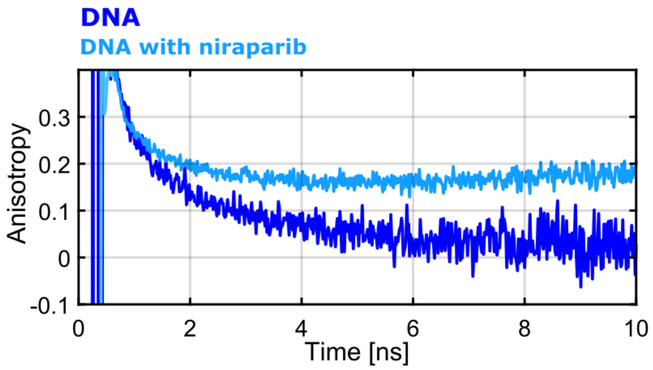


**Supplementary Figure S8: Donor anisotropy decays for DNA (blue) and DNA with Niraparib (light blue)**

The anisotropy decay (Supplementary methods) for DNA in presence of niraparib indicates a strong hindrance of the donor dye attached DNA (residual anisotropy $r_\infty$ of DNA with niraparib is higher than that of DNA in absence of niraparib), whereas the donor dye without niraparib rotates freely ($r_\infty$ approaches 0). This hindrance of the donor dye results in slightly lower FRET efficiencies for DNA with PARP-1 and niraparib data (Fig. 6 B, turquoise, Supplementary Fig. S2, Supp. Table S1).

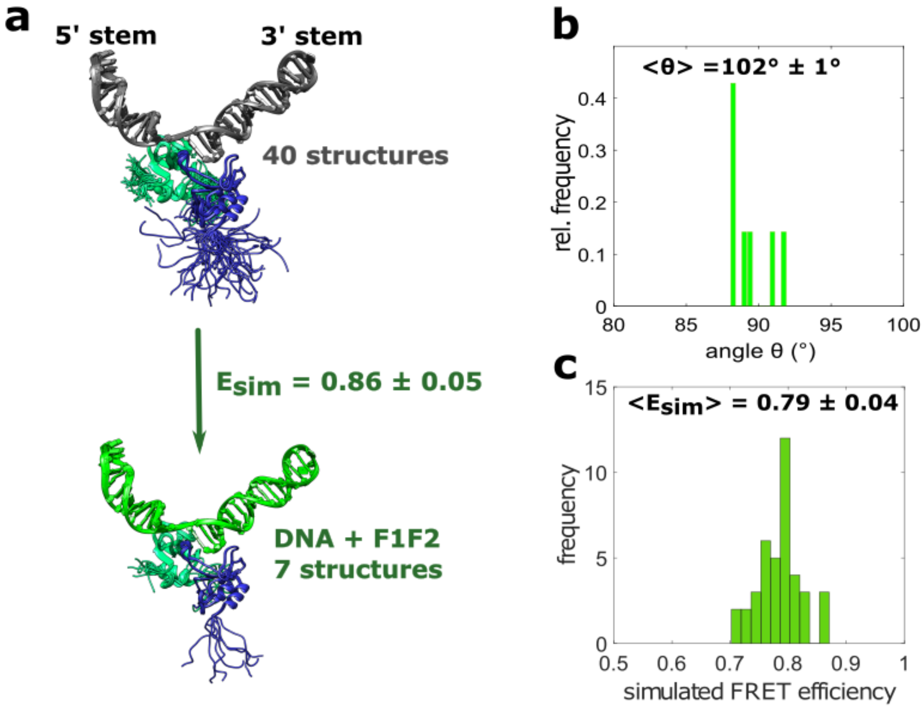


**Supplementary Figure S9: Simulated structure of nicked DNA bound by F1F2.**

**a:** Ensemble of 40 model structures for the DNA ligand bound to F1F2. Both stems of the DNA are modeled in B-form. Contacts between F1 and the 5' stem, between F2 and the 3' stem and between the two zinc fingers were modeled, and as well orientational restraints from residual dipolar coupling data were included in the calculations of these models, using values measured in the context of a similar (1nt gapped) DNA ligand by NMR (Eustermann et al. 2015). The kinking angle of the DNA is defined as the angle between the axes of the two stems. DNA in grey, F1 in light green, F2 in purple, zinc ions in brown. **b:** Histogram of kinking angles for the structures of the DNA bound to F1F2 shown in A. The mean and standard deviation of the angle distribution is shown. **c:** Histogram of simulated FRET efficiencies for the 40 structures of the DNA bound to F1F2 shown in A. Mean and standard deviation are given above the plot.

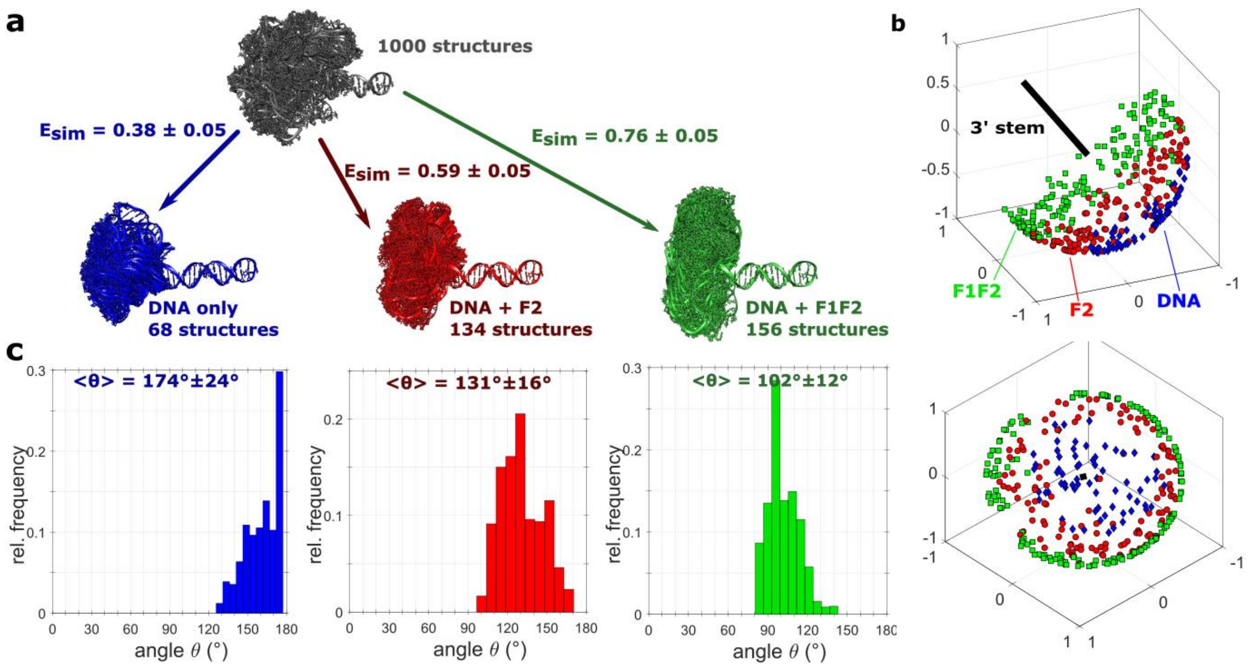


**Supplementary Figure S10: Selection and analysis of structures in accordance with experiment for the alternative DNA construct with TAMRA as donor ($DNA_{Tamra}$).**

**a:** Structures in agreement with the experimental FRET efficiencies (see Fig. 2) were selected from the ensemble of DNA (1000 structures, 100 shown in grey): *E*=0.38 for free DNA (68 structures in blue), *E*=0.59 for DNA+F2 (134 structures in red), *E*=0.76 for DNA+F1F2 (156 structures in green). An uncertainty of ΔE=±0.05 was considered in all cases. **b:** Combined representation of the selected structures shown in a. All structures are aligned with respect to the 3' stem. The axis of the 3' stem is represented by a black line. The colored shapes indicate the direction of the 5' axis for each selected structure: blue diamonds for free DNA; red circles for DNA+F2; green squares for DNA+F1F2. Two different views of the 3D plot are shown, a side view (top) and a view along the 3' axis (bottom). **c:** Kinking angle distributions for the selected structures in a. From left to right: free DNA (blue), DNA+F2 (red), DNA+F1F2 (green). The mean and standard deviation of the angles are indicated for each histogram (Kallis 2020).

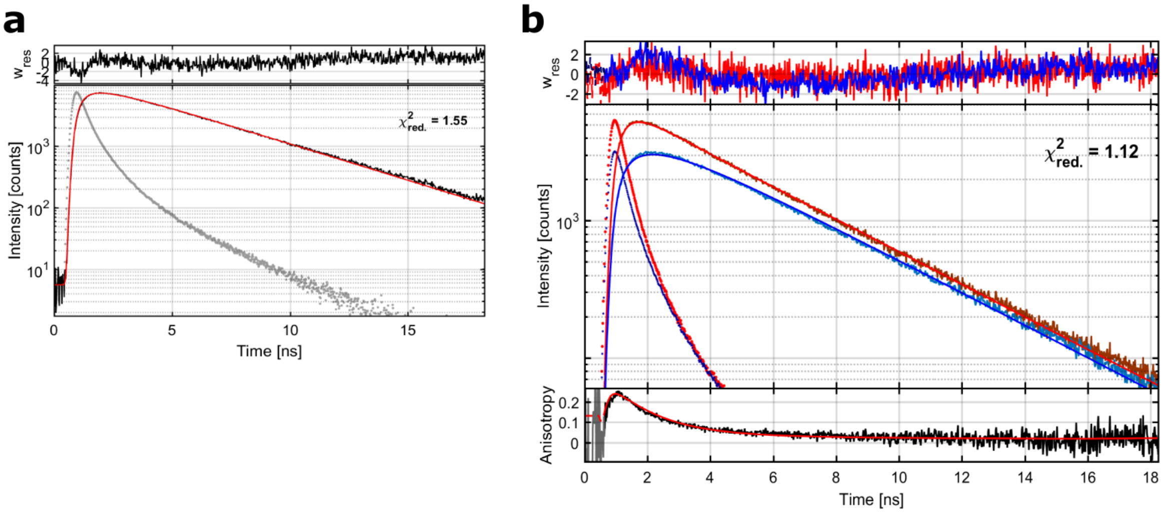


**Supplementary Figure S11: Exemplary lifetime and anisotropy fits and fit results for $DNA_{Atto}$ in presence of F1F2.**

Lifetime and anisotropy analysis (Supplementary methods) for DNA, DNA+F2 and DNA+F1F2 was performed to obtain the parameters (Supp. Table S4) necessary for the structural analysis (Methods). **a:** Fitted (red) donor fluorescence decay histogram (black), instrument response function (grey); the weighted residuals ($w_{res}$) are shown in the top panel of the fluorescence decay. The fit returned a value of $\tau_1 = 3.52n$. **b:** Fitted donor time-resolved polarized fluorescence decays (red is the parallel channel, blue is the perpendicular channel) and the dotted decays correspond to respective IRFs; the lower panel shows the anisotropy decay curve (black) and its fit function (red); the weighted residuals ($w_{res}$) for both channels are shown in the top panel of the fluorescence decay. The parameters returned from the fit are $r_0 = 0.27n$, $r_\infty = 0.04ns$, $\rho_1 = 1.31$, for which the lifetime was fixed to the values given in a.

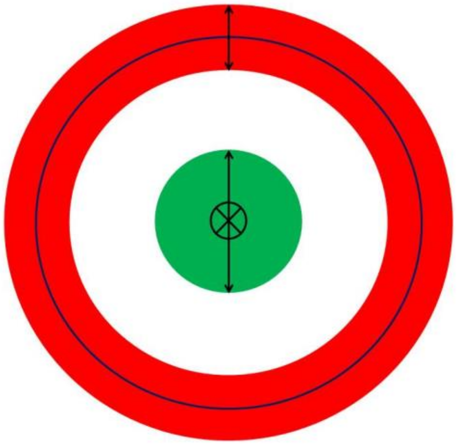

**Supplementary Figure S12:** Top view on a spherical cone (green) and an open spherical sector (red). Both are directed along the same axis here, indicated by the black crossed circle (center). This axis is also the mean axis of the cone, which does not need to coincide with the 3'-stem. However, we assume that the open spherical sector is directed along the 3'-axis. Its degenerate mean orientation is here shown by the black ring. The standard deviations along the spherical cone or the open spherical sector are indicated by a black double arrows.

**Supplementary Table S1:** Results of Gaussian fitting of smFRET efficiency histograms in Supplementary Fig. S2. *A*: area fraction, *E*: mean FRET efficiency, $\sigma$: standard deviation of the Gaussian. FRET efficiencies of relevant species are highlighted according to the color code in Supplementary Fig.S2.

| | $A_1$ | $E_1$ | $\sigma_1$ | $A_2$ | $E_2$ | $\sigma_2$ | $A_3$ | $E_3$ | $\sigma_2$ | $R^2$ |
|---|---|---|---|---|---|---|---|---|---|---|
| **DNA** | 0.77 | 0.499 | 0.096 | 0.23 | 0.39 | 0.141 | | | | 0.81 |
| **ligated DNA** | 0.75 | 0.484 | 0.092 | 0.25 | 0.39 | 0.126 | | | | 1.74 |
| **DNA with F2** | 0.22 | 0.69 | 0.567 | 0.19 | 0.49 | 0.155 | 0.56 | 0.61 | 0.117 | 1.89 |
| **DNA with F1F2** | 0.44 | 0.86 | 0.039 | 0.30 | 0.58 | 0.144 | 0.25 | 0.77 | 0.056 | 2.78 |
| **DNA with PARP-1** | 0.35 | 0.87 | 0.040 | 0.35 | 0.53 | 0.157 | 0.29 | 0.78 | 0.066 | 1.87 |
| **DNA with PARP-1 and Eb-47** | 0.45 | 0.87 | 0.038 | 0.19 | 0.57 | 0.154 | 0.36 | 0.78 | 0.067 | 2.50 |
| **DNA with PARP-1 and Niraparib** | 0.32 | 0.81 | 0.062 | 0.46 | 0.38 | 0.142 | 0.22 | 0.69 | 0.086 | 1.67 |

**Supplementary Table S2:** fFCS fit results for DNA with F2, F1F2 and PARP-1. For each dataset two auto- (Species 1 x Species 1 and Species 2 x Species 2) and two cross-correlation functions (Species 1 x Species 2 and Species 2 x Species 1) were calculated and then fitted (Methods). From the fit, the parameters shown in the table were returned: N: mean number of molecules in the confocal volume, $\tau_D$: diffusion time, $\tau$: relaxation time with the respective fraction A and $\tau_{phot}$: relaxation time due to acceptor photophysics and its respective fraction $A_{phot}$, $y_0$: offset and the weighted residuals $\chi^2$ indicating the goodness of the fit. The blue shaded values were fixed according to previous FCS experiments (Supplementary Fig. S6) and the gray shaded value results from a global fit for the different correlations.

| **Data set** | Correlation | N | $\tau_D$[μs] | $\tau$[μs] | A | $\tau_{phot}$[μs] | $A_{phot}$ | $y_0$ | $\chi^2$ |
|---|---|---|---|---|---|---|---|---|---|
| **DNA with F2** | **Sp1 x Sp1** | 0.036 | 1411 | 404 | 0.44 | 7.01 | 0.37 | -0.64 | 3.19 |
| | **Sp1 x Sp2** | 0.040 | | | 0.27 | | 0.63 | -0.58 | 2.47 |
| | **Sp2 x Sp1** | 0.041 | | | 0.29 | | 0.52 | -0.50 | 2.19 |
| | **Sp2 x Sp2** | 0.016 | | | 0.20 | | 0.59 | -1.67 | 2.40 |
| **DNA with F1F2** | **Sp1 x Sp1** | 0.029 | 1587 | 265 | 0.40 | 7.38 | 0.21 | -0.71 | 1.90 |
| | **Sp1 x Sp2** | 0.075 | | | 0.44 | | 0.37 | -0.40 | 1.28 |
| | **Sp2 x Sp1** | 0.075 | | | 0.44 | | 0.37 | -0.27 | 1.32 |
| | **Sp2 x Sp2** | 0.019 | | | 0.09 | | 0.28 | -1.4 | 1.91 |
| **DNA with PARP-1** | **Sp1 x Sp1** | 0.042 | 2342 | 273 | 0.43 | 9.58 | 0.22 | -0.65 | 0.87 |
| | **Sp1 x Sp2** | 0.099 | | | 0.37 | | 0.43 | -0.53 | 1.34 |
| | **Sp2 x Sp1** | 0.100 | | | 0.38 | | 0.47 | -0.39 | 1.22 |
| | **Sp2 x Sp2** | 0.018 | | | 0.053 | | 0.31 | -1.88 | 1.44 |

**Supplementary Table S3:** Protein Sequence of the PARP-1 fragments

| F2, 103-214 |
| --- |
| GSKAEKTLGDFAAEYAKSNRSTCKGCMEKIEKGQVRLSKKMVDPEKPQLGMIDRWYHPGCFVKNREELGFRPEYSA SQLKGFSLLATEDKEALKKQLPGVKSEGKRKGDEVD |
| **F1F2, 1-214** |
| MAESSDKLYRVEYAKSGRASCKKCSESIPKDSLRMAIMVQSPMFDGKVPHWYHFSCFWKVGHSIRHPDVEVDGFSE LRWDDQQKVKKTAEAGGVTGKGQDGIGSKAEKTLGDFAAEYAKSNRSTCKGCMEKIEKGQVRLSKKMVDPEKPQL GMIDRWYHPGCFVKNREELGFRPEYSASQLKGFSLLATEDKEALKKQLPGVKSEGKRKGDEVD |
| **PARP-1 fl, 1-1014** |
| MAESSDKLYRVEYAKSGRASCKKCSESIPKDSLRMAIMVQSPMFDGKVPHWYHFSCFWKVGHSIRHPDVEVDGFSE LRWDDQQKVKKTAEAGGVTGKGQDGIGSKAEKTLGDFAAEYAKSNRSTCKGCMEKIEKGQVRLSKKMVDPEKPQL GMIDRWYHPGCFVKNREELGFRPEYSASQLKGFSLLATEDKEALKKQLPGVKSEGKRKGDEVDGVDEVAKKKSKKEK DKDSKLEKALKAQNDLIWNIKDELKKVCSTNDLKELLIFNKQQVPSGESAILDRVADGMVFGALLPCEECSGQLVFKS DAYYCTGDVTAWTKCMVKTQTPNRKEWVTPKEFREISYLKKLKVKKQDRIFPPETSASVAATPPPSTASAPAAVNSSA SADKPLSNMKILTLGKLSRNKDEVKAMIEKLGGKLTGTANKASLCISTKKEVEKMNKKMEEVKEANIRVVSEDFLQDV SASTKSLQELFLAHILSPWGAEVKAEPVEVVAPRGKSGAALSKKSKGQVKEEGINKSEKRMKLTLKGGAAVDPDSGLE HSAHVLEKGGKVFSATLGLVDIVKGTNSYYKLQLLEDDKENRYWIFRSWGRVGTVIGSNKLEQMPSKEDAIEHFMKL YEEKTGNAWHSKNFTKYPKKFYPLEIDYGQDEEAVKKLTVNPGTKSKLPKPVQDLIKMIFDVESMKKAMVEYEIDLQK MPLGKLSKRQIQAAYSILSEVQQAVSQGSSDSQILDLSNRFYTLIPHDFGMKKPPLLNNADSVQAKVEMLDNLLDIEV AYSLLRGGSDDSSKDPIDVNYEKLKTDIKVVDRDSEEAEIIRKYVKNTHATTHNAYDLEVIDIFKIEREGECQRYKPFKQL HNRRLLWHGSRTTNFAGILSQGLRIAPPEAPVTGYMFGKGIYFADMVSKSANYCHTSQGDPIGLILLGEVALGNMYE LKHASHISKLPKGKHSVKGLGKTTPDPSANISLDGVDVPLGTGISSGVNDTSLLYNEYIVYDIAQVNLKYLLKLKFNFKTS LW |

**Supplementary Table S4:** Input parameters for calculation of accessible volumes. Atom C7 of thymidine T37 or T18 was used as attachment point for donor and acceptor, respectively. For the acceptor Alexa647, the diameter of 12Å represents a balance between the different dimensions of the dye, as it has a rather elongated structure.

| | | $DNA_{Atto}$ | | $DNA_{Tamra}$ | |
| --- | --- | --- | --- | --- | --- |
| | | **donor** | **acceptor** | **donor** | **acceptor** |
| dye diameter | **d [Å]** | 12 | 12 | 12 | 12 |
| linker length | **l [Å]** | 13 | 13 | 13 | 18 |
| linker width | **w [Å]** | 4.5 | 4.5 | 4.5 | 4.5 |
| skel. dist. | **s [Å]** | 2 | | 2 | |

**Supplementary Table S5:** Input parameters for simulation of FRET efficiencies.

| | | $DNA_{Atto}$ | $DNA_{Tamra}$ |
| --- | --- | --- | --- |
| Förster radius | **$R_0$ [Å]** | 70 | 68 |
| lifetime D | **$\tau_D$ [ns]** | 3.53 | 2.57 |
| rot. corr. time D | **$\rho_D$ [ns]** | 1.34 | 0.79 |
| residual aniso. D | **$r_{\infty D}$** | 0.042 | 0.153 |
| rot. corr. time A | **$\rho_A$ [ns]** | 0.70 | 0.34 |
| residual aniso. A | **$r_{\infty A}$** | 0.126 | 0.154 |
| number of excitations | **trajN** | 50000 | 50000 |

Kallis, Eleni. 2020. "Quantitative Single-Molecule FRET and Its Application in Experiments on DNA Damage Recognition by PARP-1." *PhD Disseration*.

Ries, Jonas, and Petra Schwille. 2012. "Fluorescence Correlation Spectroscopy." *BioEssays* 34 (5): 361–68. https://doi.org/10.1002/bies.201100111.

Schwille, Petra, F.J. Meyer-Almes, and Rudolf Rigler. 1997. "Dual-Color Fluorescence Cross-Correlation Spectroscopy for Multicomponent Diffusional Analysis in Solution." *Biophysical Journal* 72 (4): 1878–86. https://doi.org/10.1016/S0006-3495(97)78833-7.

Stennett, Elana M. S., Monika A. Ciuba, Su Lin, and Marcia Levitus. 2015. "Demystifying PIFE: The Photophysics Behind the Protein-Induced Fluorescence Enhancement Phenomenon in Cy3." *The Journal of Physical Chemistry Letters* 6 (10): 1819–23. https://doi.org/10.1021/acs.jpclett.5b00613.

Taniguchi, Masahiko, Hai Du, and Jonathan S. Lindsey. 2018. "PhotochemCAD 3: Diverse Modules for Photophysical Calculations with Multiple Spectral Databases." *Photochemistry and Photobiology* 94 (2): 277–89. https://doi.org/10.1111/php.12862.

Torella, Joseph P., Seamus J. Holden, Yusdi Santoso, Johannes Hohlbein, and Achillefs N. Kapanidis. 2011. "Identifying Molecular Dynamics in Single-Molecule Fret Experiments with Burst Variance Analysis." *Biophysical Journal* 100 (6): 1568–77. https://doi.org/10.1016/j.bpj.2011.01.066.

Torres, Tedman, and Marcia Levitus. 2007. "Measuring Conformational Dynamics: A New FCS-FRET Approach." *Journal of Physical Chemistry B* 111 (25): 7392–7400. https://doi.org/10.1021/jp070659s.
